# Disease-linked TDP-43 hyperphosphorylation suppresses TDP-43 condensation and aggregation

**DOI:** 10.1101/2021.04.30.442163

**Authors:** Lara Gruijs da Silva, Francesca Simonetti, Saskia Hutten, Henrick Riemenschneider, Erin L. Sternburg, Lisa M. Pietrek, Jakob Gebel, Volker Dötsch, Dieter Edbauer, Gerhard Hummer, Lukas S. Stelzl, Dorothee Dormann

**Affiliations:** Johannes Gutenberg-Universität (JGU), Biocenter, Institute of Molecular Physiology, 55128 Mainz, Germany; Graduate School of Systemic Neurosciences (GSN), 82152 Planegg-Martinsried, Germany; German Center for Neurodegenerative Diseases (DZNE), 81377 Munich, Germany; Max Planck Institute of Biophysics, Department of Theoretical Biophysics, 60438 Frankfurt am Main, Germany; Institute for Biophysical Chemistry, Goethe-Universität, 60438 Frankfurt am Main, Germany; Munich Cluster for Systems Neurology (SyNergy) Munich, 81377 Munich, Germany; Institute for Biophysics, Goethe-Universität, 60438 Frankfurt am Main, Germany; Johannes Gutenberg-Universität (JGU), KOMET1, Institute of Physics, 55128 Mainz, Germany; Institute of Molecular Biology (IMB), 55128 Mainz, Germany

**Keywords:** Neurodegeneration, phase separation, phosphorylation, RNA-binding protein, TDP-43

## Abstract

Post-translational modifications (PTMs) have emerged as key modulators of protein phase separation and have been linked to protein aggregation in neurodegenerative disorders. The major aggregating protein in amyotrophic lateral sclerosis (ALS) and frontotemporal dementia (FTD), the RNA-binding protein TDP-43, is hyperphosphorylated in disease on several C-terminal serine residues, which is generally believed to promote TDP-43 aggregation. Here, we show that hyperphosphorylation by Casein kinase 1δ or C-terminal phosphomimetic mutations surprisingly reduce TDP-43 phase separation and aggregation and render TDP-43 condensates more liquid-like and dynamic. Multi-scale simulations reveal reduced homotypic interactions of TDP-43 low complexity domains through enhanced solvation of phosphomimetic residues. Cellular experiments show that phosphomimetic substitutions do not affect nuclear import or RNA regulatory functions of TDP-43, but suppress accumulation of TDP-43 in membrane-less organelles and promote its solubility in neurons. We propose that TDP-43 hyperphosphorylation may be a protective cellular response to counteract TDP-43 aggregation.

## Introduction

TAR DNA binding protein (TDP-43) is the major aggregating protein in ALS and FTD patients and also forms pathological aggregates in up to 50% of Alzheimer’s disease patients (Josephs et al., 2014; Neumann et al., 2006). It is a ubiquitously expressed RNA-binding protein (RBP) with key functions in RNA processing, e.g. regulation of alternative splicing and polyadenylation, miRNA processing, mRNA stability and localization (Ratti and Buratti, 2016). In the affected brain regions of ALS and FTD patients, the physiological diffuse nuclear localization of TDP-43 is lost. Instead the protein forms cytoplasmic and occasionally nuclear inclusions in neurons and glial cells (Mackenzie et al., 2010). TDP-43 pathology closely correlates with neurodegeneration, and both loss-of-function mechanisms, e.g. misregulation of nuclear RNA targets, and gain-of-function mechanisms, e.g. aberrant interactions of the TDP-43 aggregates, are believed to contribute to neuronal dysfunction and eventually neurodegeneration (Ling et al., 2013; Tziortzouda et al., 2021).

Like other prion-like RBPs, TDP-43 is thought to aggregate through aberrant liquid-liquid phase separation (LLPS), i.e. the transition of liquid-like RBP condensates into a solid-like state (Nedelsky and Taylor, 2019). Aberrant phase transitions may occur in stress granules (SGs) or other membrane-less organelles (MLOs), where aggregation-prone RBPs are highly concentrated and exceed the critical concentration for LLPS (Alberti and Dormann, 2019; Alberti and Hyman, 2021). Subsequent liquid-to-solid phase transition, as demonstrated for various disease-linked RBPs *in vitro* (Molliex et al., 2015; Patel et al., 2015), may then cause formation of pathological RBP inclusions. LLPS is often driven by intrinsically disordered low complexity domains (LCDs), that tend to engage in weak multivalent interactions with other molecules (Alberti, 2017). TDP-43 harbors a long C-terminal LCD enriched in glycine, serine, asparagine and glutamine residues, which drives intermolecular TDP-43 interactions and assembly by phase separation (Babinchak et al., 2019; Conicella et al., 2016). The LCD is also the region that harbors numerous ALS-linked point mutations (Buratti, 2015), suggesting that small chemical changes to the TDP-43 LCD can cause neurodegeneration.

LLPS and MLO dynamics are often regulated by post-translational modifications (PTMs) in LCDs, as the introduction of small chemical groups or proteins changes the chemical nature of amino acids, e.g. their charge or hydrophobicity, which can alter their molecular interactions and LLPS behavior (Bah and Forman-Kay, 2016; Hofweber and Dormann, 2019). A highly disease-specific PTM on deposited TDP-43 inclusions is hyperphosphorylation on C-terminal serine residues in the LCD (Hasegawa et al., 2008; Inukai et al., 2008; Kametani et al., 2016; Neumann et al., 2009). Antibodies specific for C-terminal TDP-43 phosphorylation sites (e.g. S409/S410 and S403/S404) detect inclusion pathology in patients, without cross-reactivity with physiological nuclear TDP-43. Therefore, C-terminal TDP-43 hyperphosphorylation is considered a pathological hallmark and is generally believed to promote TDP-43 aggregation (Buratti, 2018). This view is largely based on the observations that C-terminal TDP-43 phosphorylation correlates with inclusion pathology and that overexpression of kinases that can phosphorylate TDP-43 enhance TDP-43 aggregation and neurotoxicity (Choksi et al., 2014; Liachko et al., 2014; Nonaka et al., 2016; Taylor et al., 2018). Based on these studies, inhibition of TDP-43 phosphorylation by specific kinase inhibitors has even been proposed as a potential therapeutic strategy for ALS (Liachko et al., 2013; Martinez-Gonzalez et al., 2020; Salado et al., 2014). However, the molecular consequences of this disease-linked PTM are still poorly understood, and its effects on TDP-43 LLPS and aggregation are still unknown.

Using *in vitro, in silico* and cellular experiments, we now demonstrate that disease-linked C-terminal hyperphosphorylation of TDP-43 suppresses TDP-43 condensation and insolubility. We show this through a) *in vitro* phase separation and aggregation assays with recombinant, full-length TDP-43; b) coarse-grained and atomistic molecular dynamics simulations of condensates of TDP-43 LCDs, elucidating molecular driving forces; and c) experiments in HeLa cells and primary rat neurons, where C-terminal phosphomimetic mutations do not disturb nuclear import or RNA processing functions of TDP-43, but abrogate TDP-43 condensation into stress-induced membrane-less organelles and enhance its solubility. Based on our findings, we suggest that C-terminal TDP-43 hyperphosphorylation may be a protective cellular response to counteract TDP-43 solidification, rather than being a driver of TDP-43 pathology, as has so far been assumed.

## Results

### *In vitro* phosphorylation with Casein kinase 1δ reduces condensation of TDP-43

To examine how phosphorylation affects TDP-43 phase transitions, we expressed and purified unphosphorylated full-length TDP-43 with a solubilizing MBP tag and a His_6_-tag in *E. coli* (Wang et al., 2018) (Fig. S1A-E). We then *in vitro* phosphorylated the purified protein with casein kinase 1 delta (CK1δ), a kinase previously reported to phosphorylate TDP-43 at disease-associated sites (Kametani et al., 2009), and confirmed phosphorylation of C-terminal serines (S403/S404; S409/S410) with phospho-specific antibodies (Fig. S2A). Mass spectrometric analysis detected phosphorylation on several additional serine/threonine sites (Fig. S2B), and the running behavior in SDS-PAGE suggests hyperphosphorylation on multiple sites (Fig. S2A, Fig. 1B). We then induced phase separation of the unphosphorylated vs. *in vitro* phosphorylated TDP-43 by cleaving off the MBP tag with TEV protease (Wang et al., 2018) and used centrifugation to separate the condensates (C) from the cleared supernatant (S) (Fig. 1A). Cleaved TDP-43 was mostly in the condensate fraction (S/C ratio <0.5), whereas *in vitro* phosphorylated TDP-43 was predominantly in the supernatant (S/C ratio ~1.8) (Fig. 1B, C). Reduced sedimentation of TDP-43 was not seen upon addition of ATP or CK1δ alone, suggesting that it is indeed caused by the addition of phospho-groups to TDP-43.

**Figure 1.**
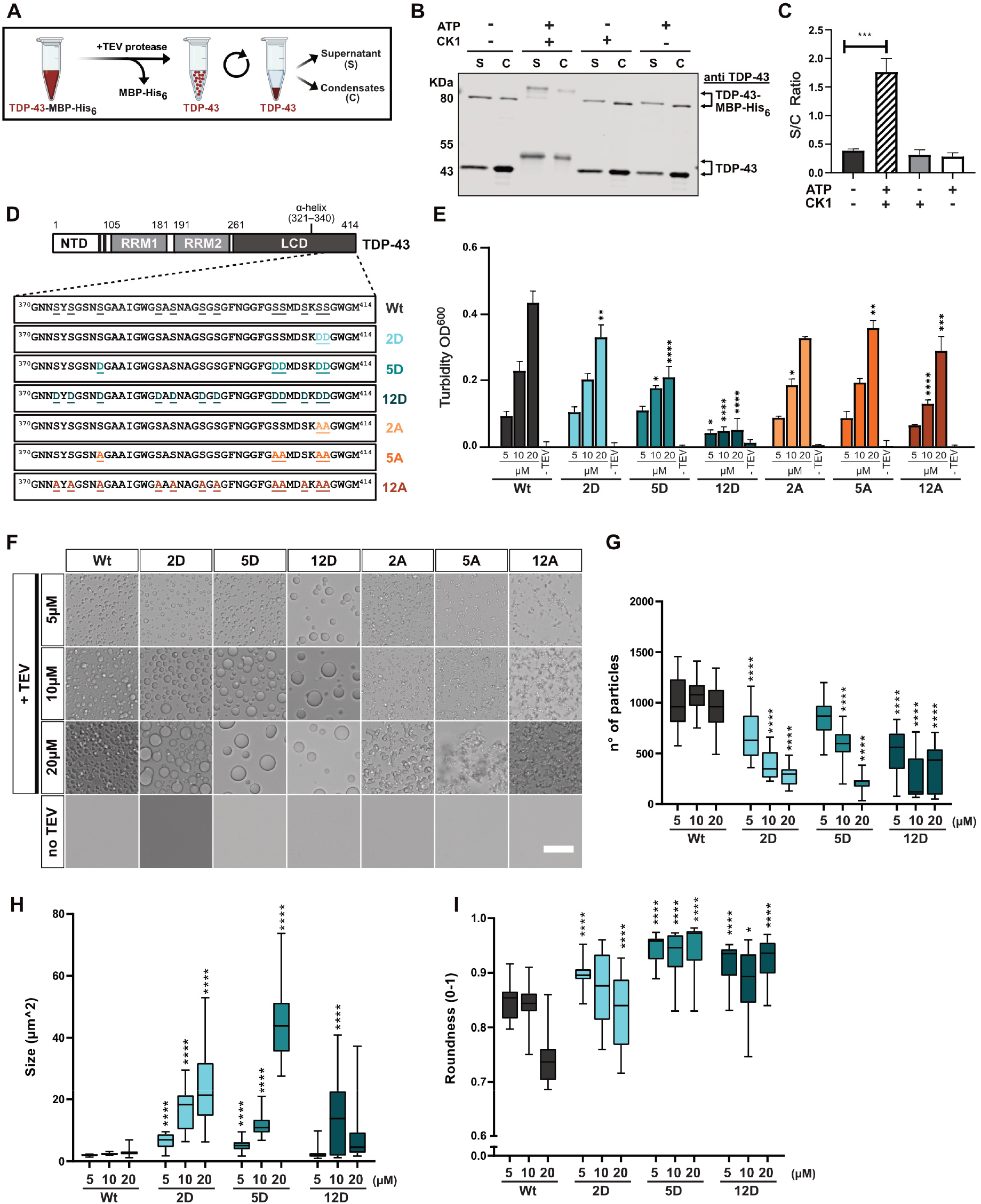
TDP-43 phosphorylation by CK1δ and C-terminal phosphomimetic substitutions reduce TDP-43 condensation *in vitro*. A Scheme of sedimentation assay (created in BioRender.com): Phase separation of TDP-43 was induced by TEV protease cleavage, and condensates were pelleted by centrifugation. B Sedimentation assay to quantify condensation of unmodified TDP-43 versus *in vitro* phosphorylated TDP-43 (+CK1δ, +ATP) and controls (CK1δ or ATP only); TDP-43 detected by Western blot (rabbit anti-TDP-43 N-term). C Quantification of band intensities shown as means of Supernatant/Condensate (S/C) ratio (n=3) ± SEM. D Schematic diagram of TDP-43 and sequence of C-terminal region for Wt, phosphomimetic (S-to-D) variants and control (S-to-A) variants. E Turbidity measurements to quantify phase separation of the indicated TDP-43 variants at three different concentrations. Values represent means (n=3) ± SD. F-I Representative bright field microscopic images of TDP-43 condensates, Bar, 25 μm (F) and quantification of condensate number (G), size (H) and roundness (I). Values represent means of all fields of view (FOV) from Min to Max (whiskers) of two replicates (≥ 22 FOV per condition). *p < 0.0332, **p < 0.0021, ***p < 0.0002 and ****p < 0.0001 by one-way ANOVA with Dunnett’s multiple comparison test to Wt (in C, E, G, H and I, respectively).

### C-terminal phosphomimetic substitutions mimicking disease-linked phosphorylation suppress TDP-43 phase separation

To study defined disease-linked phosphorylation sites, we generated phosphomimetic proteins harboring different numbers of phosphomimetic serine-to-aspartate (S-to-D) mutations (or corresponding serine-to-alanine (S-to-A) mutations as control) in the C-terminal region of TDP-43. Phosphorylation on S409/S410 is a highly specific and consistent feature of aggregated TDP-43 in all ALS/FTD subtypes (Inukai et al., 2008; Neumann et al., 2009), and five phosphorylation sites (S379, S403, S404, S409 and S410) were detected with phosphorylation site-specific antibodies in human post-mortem tissue (Hasegawa et al., 2008). Therefore, we mutated these serines to create “2D” and “5D” variants as well as the corresponding “2A” and “5A” controls (Fig. 1D). Based on a mass spectrometric study that found phosphorylation on 12 out of 14 serines in the C-terminal LCD of TDP-43 in ALS spinal cord (Kametani et al., 2016), we also mutated these 12 sites (S373, S375, S379, S387, S389, S393, S395, S403, S404, S407, S409, S410) to create “12D” or “12A” variants (Fig. 1D). Interestingly, the PLAAC web tool (http://plaac.wi.mit.edu/) that allows prediction of probable prion subsequences using a hidden-Markov model (HMM) algorithm (Lancaster et al., 2014), predicted a reduced prionlike character of the phosphomimetic 12D variant in comparison to the wild-type (Wt) and 12A protein (Fig. S3).

To study phase separation experimentally, all variants were expressed and purified as TDP-43-MBP-His_6_ fusion proteins (Fig. S1A-E), and phase separation induced by TEV protease-mediated cleavage of the MBP tag was examined by turbidity, sedimentation or microscopic condensate assays. Turbidity measurements revealed a concentration-dependent increase in phase separation for TDP-43 Wt, as expected, whereas the increase was less pronounced for the 2D and 5D variants and no concentration-dependent increase was seen for the 12D mutant (Fig. 1E). The gradual decrease in turbidity caused by the phosphomimetic mutations (Wt>2D>5D>12D) was not seen to the same extent for the corresponding S-to-A control mutations (Fig. 1E), hence suppression of phase separation is not due to the loss of serines at these positions, but can be attributed to the additional negative charges introduced by the D substitutions. Turbidity assays in phosphate buffer instead of Hepes buffer gave similar results (Fig. S4A), and sedimentation assays confirmed that TDP-43 condensation is gradually suppressed by increasing numbers of phosphomimetic mutations (Fig. S4B, C).

### Phosphomimetic S-to-D substitutions lead to rounder TDP-43 condensates, whereas S-to-A mutations cause an amorphous, aggregate-like morphology

Interestingly, bright field microscopy revealed that TDP-43 Wt formed relatively small, amorphous condensates, suggestive of solid-like material properties (Fig. 1F). In contrast, the phosphomimetic S-to-D proteins formed fewer, but much larger and rounder condensates (Fig. 1F, see G-I for quantification), suggesting a more liquid-like behavior and therefore fusion of condensates into larger droplets. Again, the observed changes were correlated with the number of phosphomimetic mutations, i.e. they were most pronounced for the 12D mutant, which formed very few, but large and perfectly circular protein droplets. (Note that these few large condensates most likely escape detection in the turbidity assay due to rapid sedimentation during the assay). In contrast, the S-to-A control variants formed numerous small, amorphous condensates and had a more irregular, aggregate-like appearance than TDP-43 Wt (Fig. 1F). This phenotype suggests that the OH groups in the respective serines influence the material properties of TDP-43 and contribute to preventing its aggregation. Similar results were obtained when the assay was carried out in phosphate buffer instead of Hepes buffer, except that 12D was unable to form any visible condensates in phosphate buffer (Fig. S4D). Together, these results demonstrate that phosphomimetic substitutions mimicking disease-linked C-terminal TDP-43 phosphorylation reduce the tendency of TDP-43 to phase separate into amorphous condensates and suggest a more dynamic, liquid-like behavior of C-terminally phosphorylated TDP-43.

### C-terminal phosphomimetic substitutions yield more liquid-like, dynamic TDP-43 condensates

To test whether the phosphomimetic mutations indeed render TDP-43 more liquid-like, we performed live imaging of Alexa488-labelled Wt, 5D and 12D condensates by spinning disc confocal microcopy. For TDP-43 Wt, no fusion events were observed over a time course of several minutes. Instead the small condensates stuck to each other like “velcro balls” (movie 1, Fig. 2A). In contrast, 5D condensates occasionally and slowly fused with each other, and 12D condensates readily fused upon contact and relaxed into perfectly round spheres, indicating a liquid droplet-like nature (movies 2 and 3, Fig. 2A). To assess the mobility of TDP-43 molecules in condensates, we performed half-bleaches of condensates and analyzed fluorescent recovery after photobleaching (FRAP) in the bleached half. In TDP-43 Wt condensates, fluorescence recovered very slowly, indicating a low mobility of TDP-43 molecules, whereas recovery was faster in 5D and even faster in 12D condensates (Fig. 2B, C), in line with an increased mobility of phosphomimetic TDP-43 compared to “unmodified” TDP-43. Taken together, phosphomimetic S-to-D substitutions in the C-terminal region enhance the liquidity of TDP-43 condensates, suggesting that phosphorylation in this region might counteract TDP-43’s tendency to form solid, irreversible aggregates.

**Figure 2.**
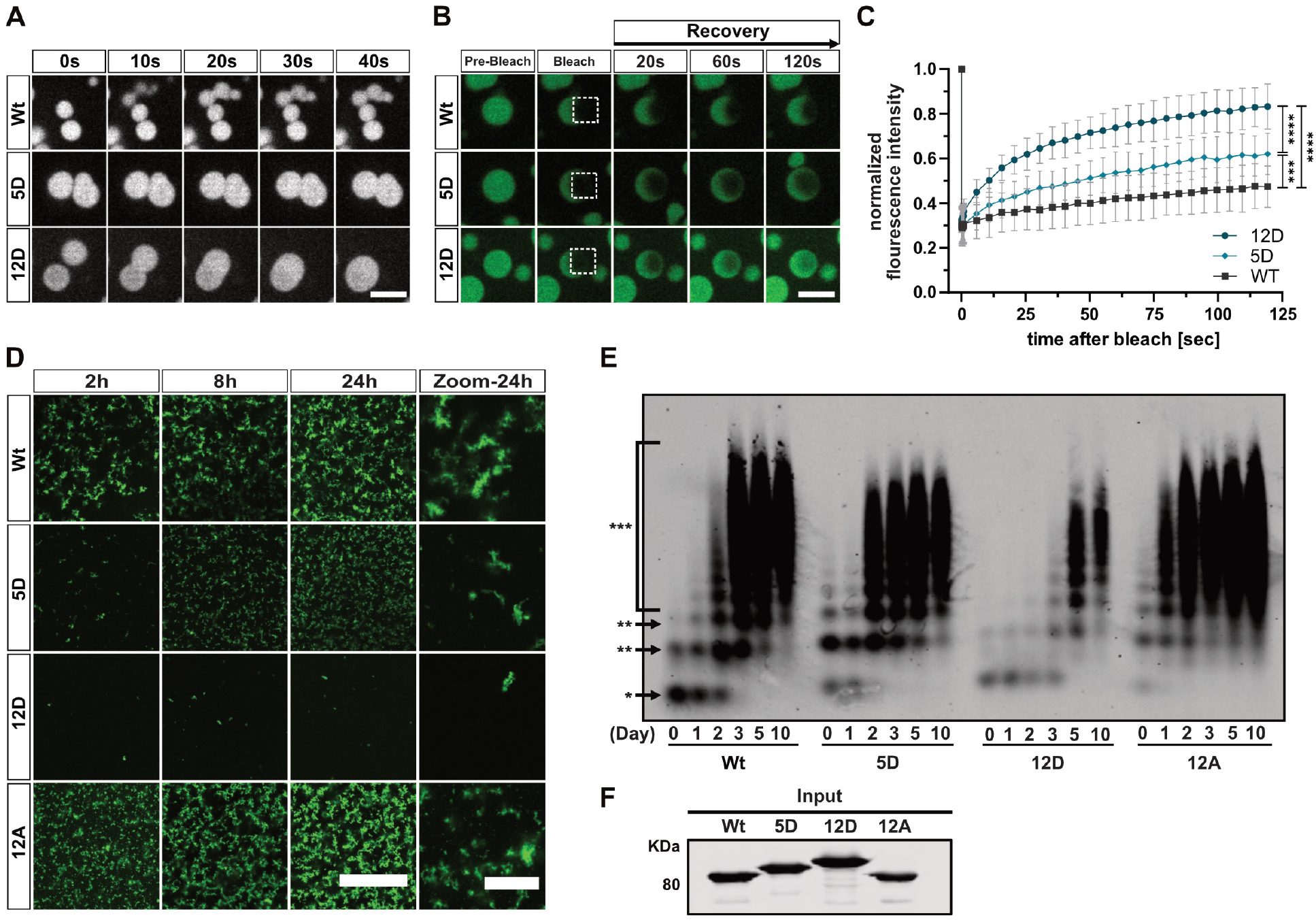
C-terminal phosphomimetic substitutions enhance liquidity of TDP-43 condensates and reduce TDP-43 aggregation *in vitro*. A Representative still images of Alexa488-labeled TDP-43 condensates by spinning disc timelapse confocal microscopy. Wt condensates do not fuse, 5D condensates fuse slowly and 12D condensates readily fuse upon contact and relax into spherical droplets. Bar, 5 μm. B Representative images of FRAP experiments at indicated time-points. Boxes indicate bleached area. Bar, 5 μm. C FRAP curves after half-bleach of freshly formed Alexa488-labeled TDP-43 condensates. Values represent means ± SD (n=3) of ≥ 9 droplets analyzed per condition. ***p < 0.0002 and ****p < 0.0001 by one-way ANOVA with Tukey’s multiple comparison test for AUC of individual droplets. D Confocal images of Alexa488-labeled TDP-43 aggregates formed in an *in vitro* aggregation assay (with TEV protease cleavage). Bar, 100 μm. Zoom shows magnified view of aggregates at the 24h time point. Bar, 20 μm. E SDD-AGE followed by TDP-43 Western blot to visualize SDS-resistant oligomers / high molecular weight species of TDP-43-MBP-His_6_ in an *in vitro* aggregation assay (without TEV protease cleavage). Asterisks represent monomeric (*), oligomeric (**) and polymeric (***) species. F Input of TDP-43 variants used in the SDD-AGE assay, detected by Western blot (anti-TDP-43 N-term).

### C-terminal phosphomimetic substitutions reduce TDP-43 aggregation

To address whether phosphorylation indeed counteracts TDP-43 aggregation, we performed *in vitro* aggregation assays modified from published protocols (French et al., 2019; Halfmann and Lindquist, 2008). Under the assay conditions, TEV cleavage of fluorescently labelled TDP-43-MBP-His_6_ yields amorphous TDP-43 aggregates that can be visualized by confocal microscopy. In contrast to Wt or 12A, the phosphomimetic 5D or 12D proteins formed much smaller and fewer aggregates, respectively (Fig. 2D), suggesting that C-terminal TDP-43 phosphorylation can efficiently suppress TDP-43 aggregation. For biochemical characterization of the formed aggregates, we performed semi-denaturing detergent-agarose gel electrophoresis (SDD-AGE) under the same assay conditions, just in the absence of TEV, as MBP-tagged TDP-43 aggregates slower than TDP-43 and distinct oligomeric/polymeric species resistant to 0.5% SDS can be visualized under these conditions (Fig. S5A). In comparison to TDP-43 Wt and 5D, 12D showed reduced and delayed oligomerization and formation of high molecular weight species (Fig. 2E, equal protein input shown in Fig. 2F). In contrast, 12A formed SDS-resistant oligomers/high molecular weight species at a higher rate, which together with our microscopic images of TDP-43 condensates (Fig. 1F), suggests that C-terminal alanine substitutions make TDP-43 more aggregation-prone. Taken together, C-terminal phosphomimetic substitutions that mimic the phosphorylation pattern in ALS patients reduce the formation of SDS-resistant high molecular weight oligomers and TDP-43 aggregates *in vitro*.

### Multiscale simulations of the TDP-43 low complexity domain reveal reduced proteinprotein interactions through enhanced solvation of phosphomimetic residues

To understand the effect of C-terminal TDP-43 phosphorylation on phase separation at the molecular level, we used coarse-grained and atomistic simulations of the disordered TDP-43 LCD (aa. 261 – 414) with and without phosphomimetic substitutions. We found that phosphomimetic substitutions locally reduce protein-protein interactions (Fig. S6) and increase protein-solvent interactions (Fig. 3). In the coarse-grained simulations, the LCD of both TDP-43 Wt and 12D phase separated spontaneously to form condensates (shown for Wt in Fig. 3A and movie 4). Yet, phosphomimicking residues are less prone to interact with protein in the phase-separated condensates and are somewhat more solvated than the corresponding serine residues (Fig. 3B, Fig. S6). The aspartate side chains in 12D LCDs engage in partially compensatory interactions with arginines, showing that introduction of charged side chains gives rise to both stabilizing and destabilizing interactions in condensates. Importantly, our simulations (Fig. S6) are in line with previous studies that have highlighted the importance of aromatic sticker-sticker interactions in driving phase separation of prion-like domains and the TDP-43 LCD (Li et al., 2018; Martin et al., 2020).

**Figure 3:**
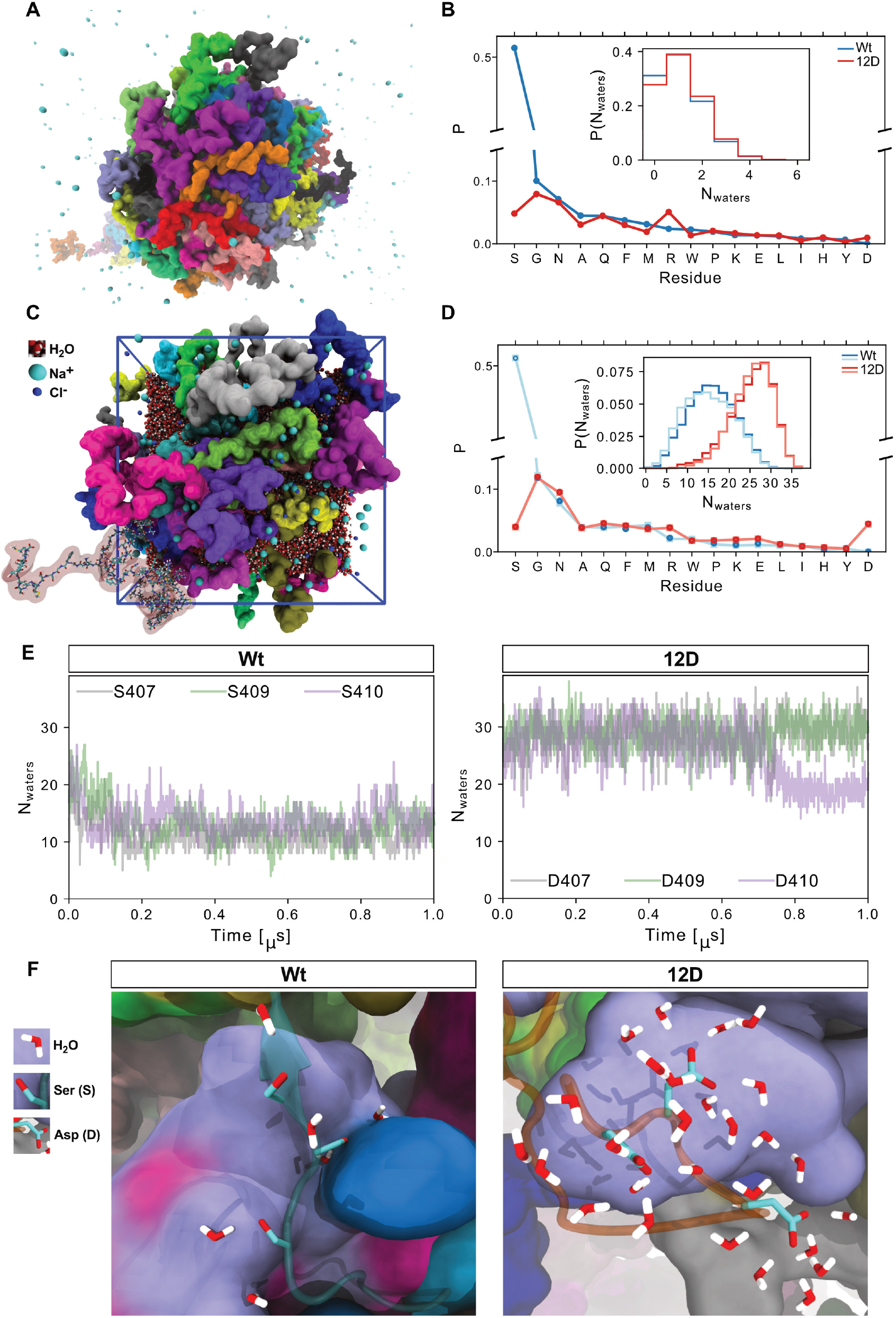
Atomistic and coarse-grained simulations of TDP-43 LCD: Phosphomimicking residues form fewer protein-protein interactions and more protein-solvent interactions. A TDP-43 LCD phase separates in coarse-grained simulations with explicit solvent. Water omitted for clarity. Ions shown in cyan. Condensate of TDP-43 Wt LCD is shown, protein colored according to chain identity. B Normalized probability of protein-protein contacts by phosphomimicking aspartates in 12D and serines in Wt resolved by amino acid type from coarse-grained simulations. Error bars smaller than symbols. Inset: Distributions of the number of water molecules within 5 Å of sidechains of phosphomimicking aspartates of 12D and corresponding serines in Wt in 3x 5 μs segments from coarse-grained simulations. Data from different segments are on top of each other. C Atomistic simulation setup of 32 TDP-43 LCDs. Different LCD chains shown in different colors in space-filling representation. For one chain (lower left), all atoms are shown to highlight the high-resolution atomistic description. D Normalized probability of protein-protein contacts by phosphomimicking aspartates in 12D and serines in Wt resolved by amino acid type from atomistic simulations. Two 1 μs simulations are distinguished by color intensity. Inset: Distributions of the number of water molecules within 5 Å of the sidechains of phosphomimicking aspartates of 12D and the corresponding serines in Wt from atomistic simulations. E Time series of number of water molecules bound to Wt S407, S409, and S410 (chain 23) and 12D D407, D409, and D410 (chain 32) in atomistic molecular dynamics. F Representative snapshots of atomistic simulations showing water within 3 Å of Wt S407, S409, S410 with nearby LCDs in surface representation and 12D D407, D409, and D410.

To characterize the interactions of TDP-43 LCDs further, we performed atomistic molecular dynamics simulations of dense protein condensates (Fig. 3C, movie 5) assembled with hierarchical chain growth (Pietrek et al., 2020) to enhance the sampling of polymeric degrees of freedom. In microsecond dynamics with explicit solvent and a highly accurate description of molecular interactions (Robustelli et al., 2018), we again found serine residues in the Wt protein to be more prone to interact with other protein residues than interacting with solvent (Fig. 3D). By contrast, phosphomimicking aspartate side chains bind more water molecules and show an overall reduced tendency for protein-protein interactions, but with some compensating interactions in particular with arginines (Fig. 3D-F, Fig. S7A). Enhanced sidechain solvation is consistent across the 12 phosphomimetic substitution sites (Fig. S7B). The simulations thus suggest that individual small effects act together to modulate the phase separation behavior of TDP-43.

### C-terminal phosphomimetic substitutions do not impair nuclear import and RNA regulatory functions of TDP-43

Next, we turned to cellular experiments to investigate how C-terminal TDP-43 phosphorylation affects the behavior and function of TDP-43 in cells. As TDP-43 hyperphosphorylation is found in the disease state, it seems possible that this PTM has detrimental effects on the protein and contributes to mislocalization and/or malfunction of TDP-43, thus driving neurodegeneration. To address this possibility, we expressed different TDP-43 variants (Wt, 12D, 12A) in HeLa cells and analyzed their intracellular localization, nuclear import and RNA processing functions. All three TDP-43 variants showed a predominantly nuclear localization (Fig. S8A), and nuclear import rates measured in a hormone-inducible reporter assay by live cell imaging (Hutten et al., 2020) were indistinguishable (Fig. 4A, B). To assess whether hyperphosphorylated TDP-43 shows functional impairments in RNA processing, we first assessed its ability to autoregulate its own levels (Avendano-Vazquez et al., 2012; Ayala et al., 2011). However, endogenous TDP-43 was downregulated to the same degree by all three TDP-43 variants (Fig. 4C), indicating that hyperphosphorylated TDP-43 can normally bind to its own 3’UTR and autoregulate its own levels. In line with these findings, recombinant TDP-43 Wt, 12D and 12A showed comparable RNA binding in electrophoretic mobility shift assays (EMSAs) with *in vitro* transcribed RNA comprised of the autoregulatory TDP-43 binding site (Fig. 4D) or synthetic (UG)_12_ RNA (Fig. S8B). Second, we examined splicing of two known TDP-43 splice targets that get mis-spliced upon loss of TDP-43 (Fiesel et al., 2012; Tollervey et al., 2011). After siRNA-mediated silencing of endogenous TDP-43 expression and reexpression of siRNA-resistant TDP-43 Wt, 12D or 12A (Fig. S8C), splicing of *SKAR* and *Bim* exon 3 were fully restored by all three TDP-43 variants (Fig. 4E), indicating normal function of phosphomimetic TDP-43 in splicing regulation. Together, our data suggest that C-terminal hyperphosphorylation is not responsible for cytosolic mislocalization or impaired RNA regulatory functions of TDP-43 in disease.

**Figure 4.**
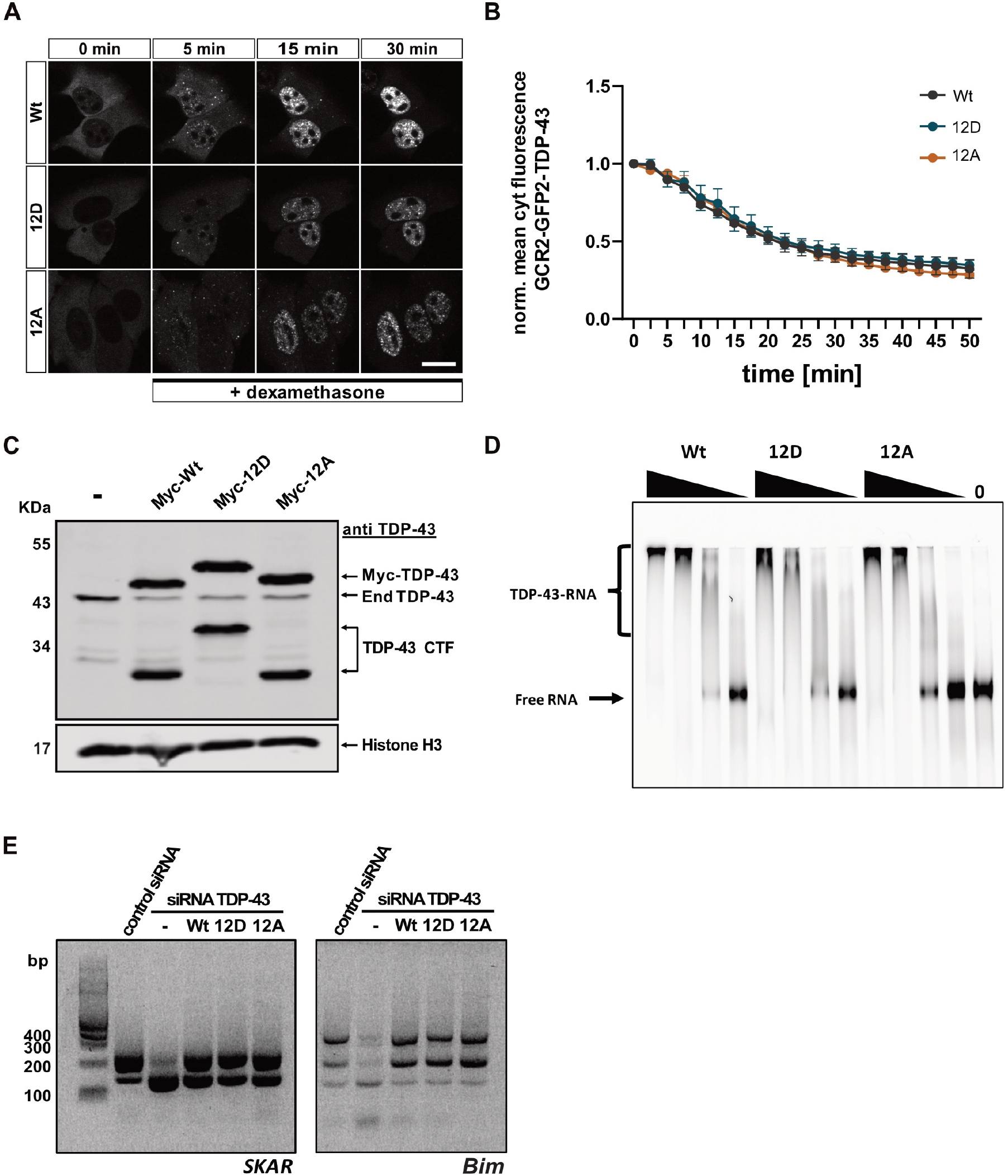
Phosphomimetic substitutions do not alter the rate of TDP-43 nuclear import and do not impair TDP-43 autoregulation, RNA-binding or alternative splicing function. A Representative still images of GCR_2_-EGFP_2_-TDP-43 Wt, 12D and 12A before and during import triggered by addition of dexamethasone. Images were live recorded by spinning disc confocal microscopy. Bar, 20 μm. B Quantification of the hormone-inducible nuclear import measured during a total time course of 50 min. Values represent the mean fluorescence intensity of TDP-43-MBP-His_6_ in the cytoplasm for three independent replicates ± SEM (≥ 42 cells per condition). C Phosphomimetic 12D TDP-43 is competent in autoregulating TDP-43 expression. SDS-PAGE followed by TDP-43 Western blot showing downregulation of endogenous TDP-43 through autoregulation (*60*) after 48 h expression of Wt, 12D and 12A variants in HeLa cells. TDP-43 was detected using rabbit anti-TDP-43 C-term antibody (Proteintech), Histone H3 (rabbit anti-Histone H3 antibody, Abcam) was visualized as a loading control. D Electrophoretic mobility shift assays (EMSA) of TDP-43-MBP-His_6_ variants (Wt, 12D and 12A) in a complex with TDP-43 autoregulatory RNA site (*60*). All TDP-43 variants form TDP-43-RNA complexes equally well. E Splicing analysis by RT-PCR of known TDP-43 splice targets (*SKAR* exon 3 and *Bim* exon 3) in HeLa cells. Silencing of endogenous TDP-43 by siRNA leads to altered splice isoforms of *SKAR* and *Bim* (second vs. first lane). These splicing alterations can be rescued by reexpression of TDP-43 Wt, but also 12D or 12A variants, demonstrating that phosphomimetic TDP-43 is fully competent in regulation splicing of these TDP-43 splice targets.

### Phosphorylation suppresses recruitment of TDP-43 into stress-induced membrane-less organelles (MLOs)

Finally, we investigated how C-terminal TDP-43 phosphorylation affects TDP-43 condensation in cellular MLOs. First, we used a quantitative assay to measure SG association of recombinant proteins under controlled conditions in semi-permeabilized HeLa cells (Hutten and Dormann, 2020) (Fig. 5A). In line with our *in vitro* condensation experiments, increasing the number of phosphomimetic S-to-D substitutions caused a gradual decrease in SG association of TDP-43 (Fig. S9A, B). *In vitro* phosphorylated TDP-43 showed a similarly strong reduction in SG association as the 12D protein (Fig. 5B, C), demonstrating that the phosphomimetic substitutions and phospho-groups introduced by a kinase have similar effects on SG association of TDP-43. Second, we expressed the different TDP-43 variants in intact HeLa cells to analyze their recruitment into stress-induced MLOs. To this end, we silenced endogenous TDP-43 expression using siRNA (Fig. S9C) and then re-introduced siRNA-resistant myc-tagged TDP-43 Wt, 12D or 12A, thus avoiding oligomerization with endogenous TDP-43 via the N-terminal domain (Afroz et al., 2017). Short term oxidative stress treatment caused a partially cytosolic relocalization of TDP-43 and led to recruitment of TDP-43 Wt and 12A, but not the 12D mutant, into TIA-1-positive SGs (Fig. 5D, E). Similar results were obtained for nuclear import-deficient TDP-43 (Fig. 5F, G) that was strongly mislocalized to the cytoplasm due to point mutations in the nuclear localization signal (NLSmut) (Fig. S9D). Finally, we examined recruitment of TDP-43 into arsenite-induced nuclear bodies (NBs) (Wang et al., 2020) and found that TDP-43 Wt and 12A were readily recruited into stress-induced NBs, while the phosphomimetic 12D protein remained dispersed in the nucleoplasm (Fig. 5H-J). Taken together, phosphomimetic substitutions that mimic disease-linked phosphorylation of TDP-43 suppress the localization of TDP-43 in phase-separated MLOs that could be condensation sites for pathological TDP-43 aggregation.

**Figure 5.**
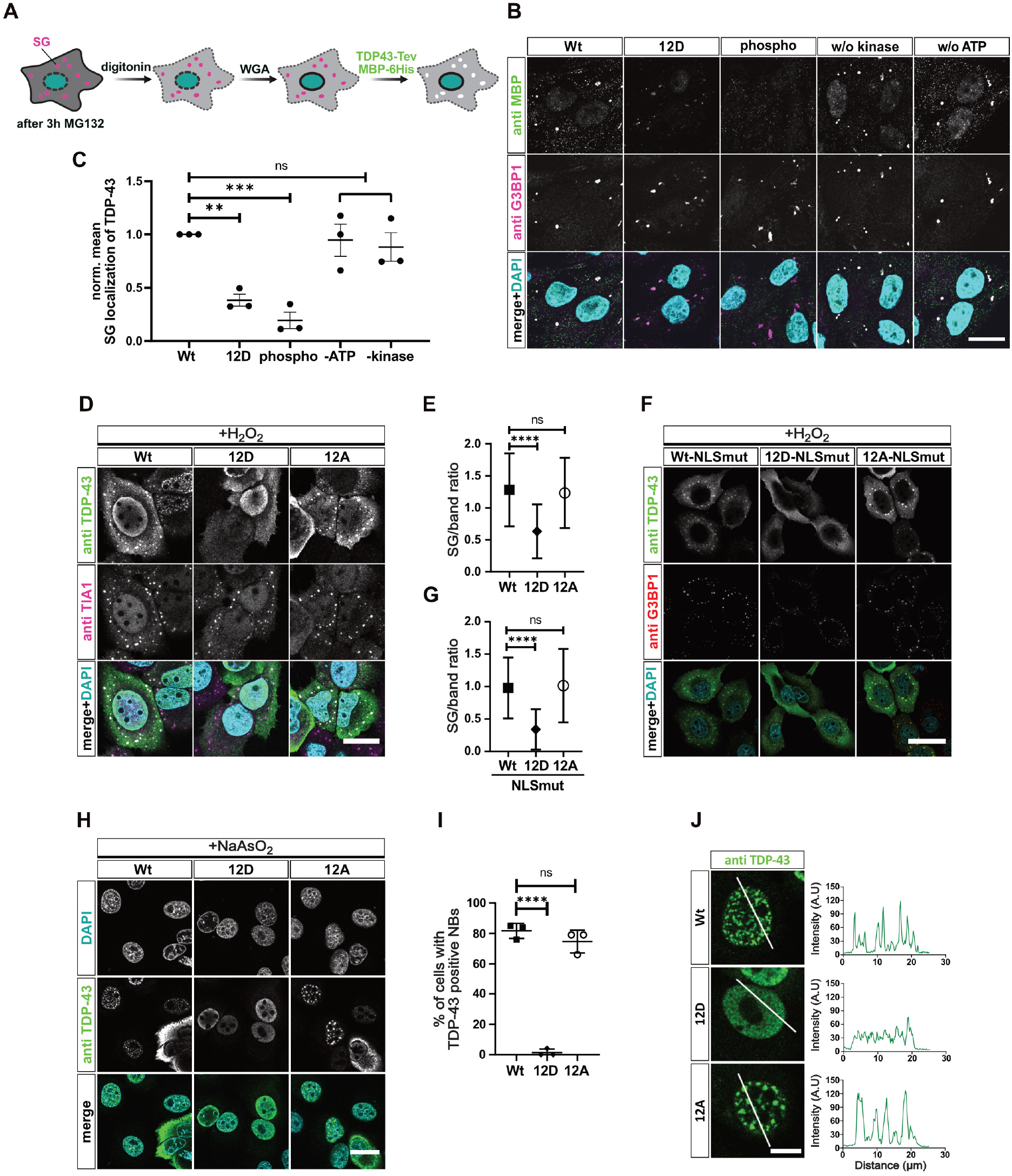
Phosphorylation and phosphomimetic substitutions reduce recruitment of TDP-43 into stress-induced membrane-less organelles. A Scheme of stress granule (SG) recruitment assay in semi-permeabilized cells. B Reduced SG association of TDP-43 by 12D mutations or *in vitro* phosphorylation. Bar, 20 μm. C Quantification of TDP-43-MBP-His_6_ mean fluorescence intensity in SGs normalized to Wt ± SEM (n=3) (≥ 10 cells; ≥ 46 SGs each). D SG recruitment of TDP-43 variants in intact HeLa cells in absence of endogenous TDP-43. After TDP-43 silencing and expression of NLSmut Wt, 12D and 12A variants, SGs were induced by H_2_O_2_ treatment and SG recruitment of TDP-43 was monitored by TDP-43 and TIA1 immunostaining. For clarity, signals were converted to grey values in the individual channels (upper two rows). In the merge (lower row), nuclei were stained in DAPI (turquoise), TDP-43 (green) and TIA-1 (magenta). Bar, 25 μm. E Quantification of TDP-43 in SGs versus cytoplasm ± SD (n=2) (≥ 62 cells; ≥ 234 SGs each). F SG recruitment of different TDP-43-NLSmut variants in intact HeLa cells in the absence of endogenous TDP-43. After TDP-43 silencing and expression of NLSmut Wt, 12D and 12A variants, SGs were induced by H_2_O_2_ treatment and SG recruitment of TDP-43 was monitored by TDP-43 and G3BP1 immunostaining. For clarity, signals were converted to grey values in the individual channels (upper two rows). In the merge (lower row), nuclei were stained in DAPI (turquoise), TDP-43 (green) and G3BP1 (red). Bar, 40 μm. G Quantification of TDP-43-NLS mutants in SGs versus band around SGs of two independent replicates ± SD. ****p < 0.0001 by one-way ANOVA with Dunnett’s multiple comparison test to Wt (≥ 56 cells; ≥ 315 SGs per condition). H Recruitment of TDP-43 into arsenite-induced nuclear bodies (NBs). Bar, 20 μm. I Percentage of cells with TDP-43 in NBs ± SD (n=3). J Intensity profiles (right) of nuclear TDP-43 Wt, 12D and 12A variants (green) along white lines (left). Bar, 10 μm.

### Phosphomimetic substitutions enhance TDP-43 solubility in Hela cells and primary neurons

To further support the idea that phosphorylation enhances the solubility of TDP-43 and counteracts its aggregation propensity in cells, we overexpressed the different TDP-43 variants in HeLa cells and performed a biochemical fractionation into a RIPA-soluble (S) and RIPA-insoluble (I) fraction. Indeed, the 12D protein had a significantly higher S/I ratio compared to the Wt and 12A proteins (Fig. 6A, B). We also expressed GFP-tagged TDP-43 Wt, 12D, 12A or the corresponding NLS-mutant cytosolic versions in primary rat neurons and then probed for RIPA-insoluble high molecular weight material in a filter trap assay. Both the nuclear and the cytosolic 12D proteins showed a strong reduction in the amount of RIPA-insoluble TDP-43 in the transduced neurons (Fig. 6C). Confocal microscopy of transduced neurons revealed a completely dispersed localization of the NLS-mutant 12D protein, whereas TDP-43 Wt and, in particular, 12A showed a more granular, condensed pattern in the neuronal cytoplasm (Fig. 6D). Thus, we conclude that phosphomimetic substitutions mimicking disease-linked C-terminal hyperphosphorylation reduce TDP-43’s tendency to condense and become insoluble in neurons and propose that TDP-43 phosphorylation might be a cellular response to counteract pathological TDP-43 aggregation.

**Figure 6.**
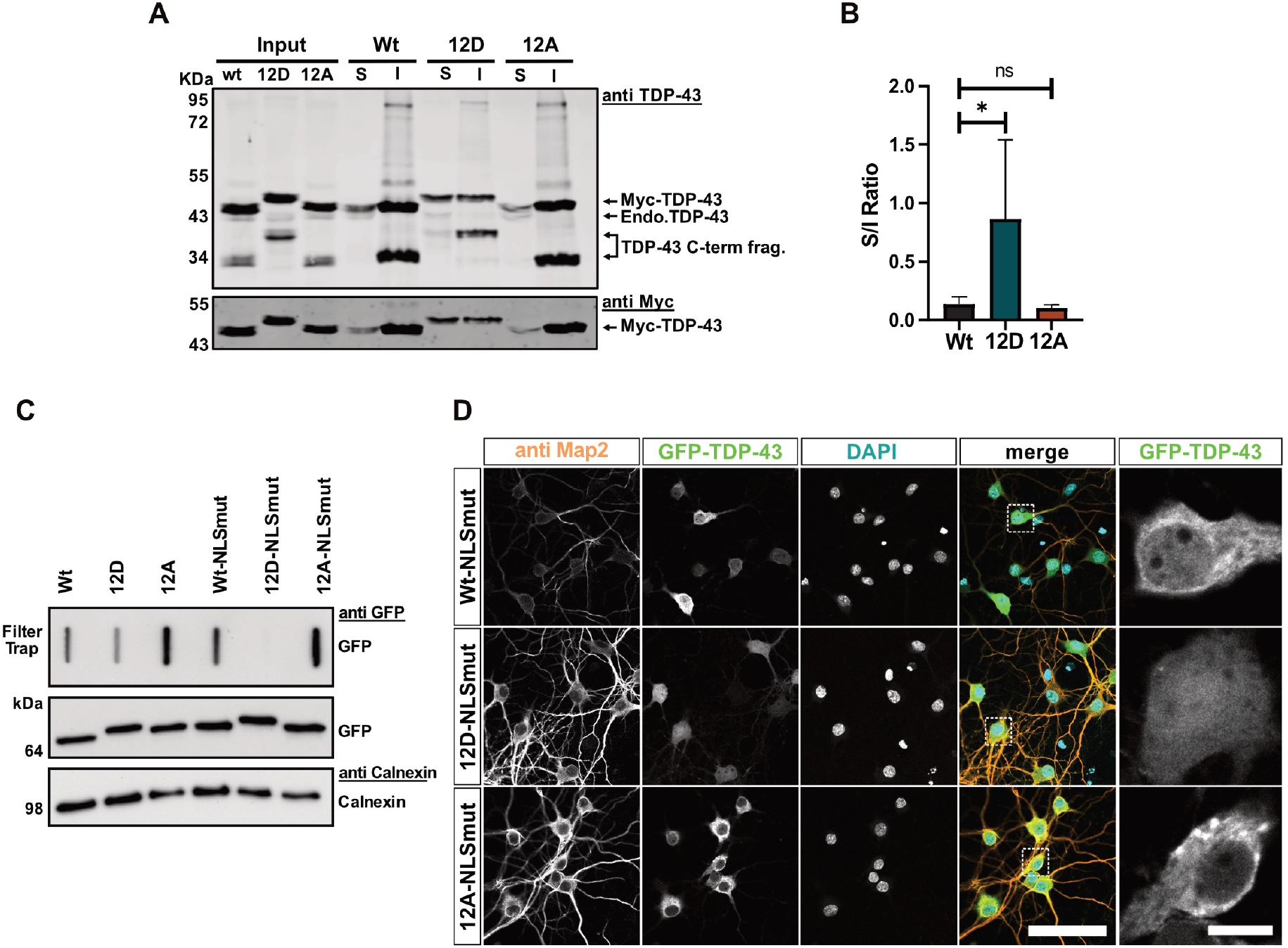
Phosphomimetic substitutions enhance TDP-43 solubility in HeLa cells and primary neurons. A Biochemical fractionation into RIPA-soluble (S) and RIPA-insoluble (I) fractions to analyze solubility of the different TDP-43 variants (Wt, 12D and 12A) expressed in HeLa cells for 48 h. TDP-43 was detected by TDP-43 Western blot (upper blot, rabbit anti-TDP-43 C-term, Proteintech) and Myc Western blot (mouse anti-Myc 9E10). B Quantification of TDP-43 variants (Wt, 12D and 12A) in (S) versus (I) fractions of 4 independent replicates ± SD. *p < 0.0332 by one-way ANOVA with Dunnett’s multiple comparison test to Wt. C RIPA-insoluble material for TDP-43 variants (± NLS mutation) in primary cortical neurons analyzed by filter-trap assay. D Primary hippocampal neurons expressing TDP-43 Wt, 12D or 12A with additional NLS mutation. Bar, 80 μm. Right: Zoomed images of white squares (TDP-43 signal). Bar, 10 μm. **p < 0.0021, ***p < 0.0002 and ****p < 0.0001 by one-way ANOVA with Dunnett’s multiple comparison test to Wt (in C, E and G).

## Discussion

C-terminal TDP-43 phosphorylation is a long-recognized pathological hallmark in ALS and FTD (Hasegawa et al., 2008; Inukai et al., 2008; Kametani et al., 2016; Neumann et al., 2009). Against previous expectations, we now show that this could be a protective rather than a pathogenic modification: We find that TDP-43 phosphorylation, and in particular phosphomimetic mutations mimicking the phosphorylation pattern in ALS/FTD (Hasegawa et al., 2008; Kametani et al., 2016), strongly suppress TDP-43 phase separation and aggregation both *in vitro* and in cells. Our data is in line with two previous studies that examined C-terminal fragments of TDP-43 with phosphomimetic 2D or 5D/E mutations and observed a reduced aggregation propensity and toxicity in cell lines and *Drosophila* (Brady et al., 2011; Li et al., 2011). Based on a recent cryo-EM structure of TDP-43 LCD fibrils, several C-terminal serines are buried inside the fibril structure (Li et al., 2021), hence phosphorylation could disrupt the amyloid fibril structure, in line with our experimental findings. We therefore propose that TDP-43 phosphorylation might be a protective cellular mechanism that counteracts aberrant TDP-43 phase transitions and renders TDP-43 more dynamic and liquid-like by reducing C-terminal LCD-LCD interactions through negatively charged, highly hydrated phospho-groups.

What triggers TDP-43 phosphorylation and how phosphorylation or the hyperphosphorylated protein is removed is still unknown. Interestingly, we and others previously found that C-terminal TDP-43 phosphorylation follows TDP-43 insolubility, suggesting that phosphorylation arises downstream of TDP-43 aggregation (Brady et al., 2011; Dormann et al., 2009). We speculate that hyperphosphorylated soluble TDP-43 is rapidly cleared, e.g. by the proteasome, and therefore is not detected under normal conditions. Under conditions of proteostasis impairment, e.g. in the aging brain and specifically in ALS/FTD patients (Hipp et al., 2019; Yerbury et al., 2020), clearance mechanisms might be impaired, leading to accumulation of hyperphosphorylated, polyubiquitinated TDP-43 that then may aggregate due to elevated protein concentration, additional PTMs, or aberrant interactions with other cellular molecules/organelles.

Several other studies on TDP-43 phosphorylation at first glance contrast our findings. Overexpression of various TDP-43 kinases in cell or animal models was shown to promote TDP-43 aggregation and neurotoxicity (Choksi et al., 2014; Liachko et al., 2014; Nonaka et al., 2016; Taylor et al., 2018). Based on these studies, inhibition of TDP-43 phosphorylation by kinase inhibitors has been proposed as a potential therapeutic strategy for ALS (Liachko et al., 2013; Martinez-Gonzalez et al., 2020; Salado et al., 2014). A possible explanation for the discrepant findings could be that kinase overexpression has pleiotropic effects that may cause TDP-43 aggregation and neurotoxicity independent of TDP-43 phosphorylation. Our data exclude such indirect effects, as they rely on experiments with purified components and defined phosphomimetic constructs rather than modulation of kinase levels/activity. Furthermore, our results suggest that beneficial effects seen with kinase inhibitors are likely not the direct consequence of reduced TDP-43 phosphorylation, but rather mediated by other mechanisms.

An alternative scenario that we cannot exclude is that reduced TDP-43 condensation due to hyperphosphorylation may have negative consequences by disturbing essential functions of TDP-43 that depend on its capacity to phase separate or solidify, e.g. certain DNA/RNA processing steps or recruitment of TDP-43 into cytoprotective NBs (Wang et al., 2020) or other MLOs. In support of this hypothesis, a deep mutagenesis study recently found that aggregating TDP-43 variants decrease toxicity in yeast, whereas dynamic, liquid-like variants enhance toxicity (Bolognesi et al., 2019), so further work is needed to investigate this possible scenario. However, our data clearly show that the core RNA processing functions (autoregulation and splicing regulation), RNA-binding and nuclear localization/import of TDP-43 are not affected by C-terminal hyperphosphorylation and therefore do not depend on TDP-43’s phase separation and solidification capacity.

Of note, abnormal PTMs are a common theme in neurodegenerative disorders, e.g. Tauopathies linked to pathological Tau aggregation (Alquezar et al., 2020; Morris et al., 2015). Interestingly, even though hyperphosphorylation is generally believed to trigger Tau aggregation, site-specific phosphorylation in the microtubule-binding region of Tau was recently shown to inhibit, rather than promote Tau fibrillization and seeding (Haj-Yahya et al., 2020). We now show that C-terminal TDP-43 phosphorylation as detected on ALS/FTD inclusions has a similar inhibitory effect on TDP-43 aggregation, underscoring the idea that aberrant PTMs detected on pathological inclusions may not necessarily all be drivers of protein aggregation, but could also have protective, anti-aggregation effects that are later-on overruled by other pathogenic mechanisms.

## Supporting information

Movie 1. Fluorescently labelled TDP-43 Wt condensates imaged live by spinning disc confocal microcopy

Movie 2. Fluorescently labelled TDP-43 5D condensates imaged live by spinning disc confocal microcopy

Movie 3. Fluorescently labelled TDP-43 12D condensates imaged live by spinning disc confocal microcopy

Movie 4. Coarse-grained simulations of TDP-43 LCD Wt with explicit solvent

Movie 5. All-atom simulations of TDP-43 LCD 12D with explicit representation of proteins, ions, and water with atomic resolution

## Acknowledgments

We thank Bettina Schmid, Sören von Bülow and Zakarya Benayad for insightful discussions and Ignasi Forné for technical support and discussion. We acknowledge support by the Core Facilities Proteomics, Bioimaging and Bioinformatics of the BMC Munich and thank Peter Becker and Michael Kiebler for infrastructure and access to the spinning disc confocal microscope (DFG, INST 86/1581-1 FUGG). This work was supported by the Deutsche Forschungsgemeinschaft (DFG, German Research Foundation) by project DO1804/4-1 within SPP2191 - ID 402723784 (to D.D.), the Munich Cluster for Systems Neurology (EXC2145 SyNergy – ID 390857198 to D.D. and D.E.) and the NOMIS foundation (D.E.). G.H. acknowledges financial support from the German Research Foundation (CRC 902: Molecular Principles of RNA Based Regulation) and the Max Planck Society. L.S.S. acknowledges support by ReALity – Resilience, Adaptation and Longevity, M3ODEL and Forschungsinitiative des Landes Rheinland-Pfalz.

## Author contributions

Conceptualization, D.D., L.G.S.; Methodology, all authors; Investigation, L.G.S., F.S., S.H., H.R., E.L.S., J.G., L.S.S.; Resources, D.D., G.H., L.S.S., D.E., V.D.; Writing – Original Draft, D. D., L.G.S.; Writing – Review and Editing, all authors; Visualization, L.G.S., F.S., S.H., H.R., E. L.S., J.G., L.S.S.; Supervision, D.D., G.H., D.E., V.D., L.S.S.; Project Administration, D.D.; Funding Acquisition, D.D., D.E., G.H., L.S.S., V.D.

## Data and materials availability

This study includes no data deposited in external repositories.

## Materials and Methods

### cDNA constructs

#### Bacterial expressing constructs

TDP-43 carrying mutations in serine 409 and 410, either to aspartate (2D) or alanine (2A) were generated by site-directed mutagenesis using Q5 high fidelity DNA polymerase (NEB) using primers containing the mutations S409D/410D and S409A/410A and pJ4M TDP-43-TEV-MBP-His_6_ vector as a template. Expression constructs with 5 or 12 serine substitutions (5D, 5A, 12D and 12A) were generated using synthetic double stranded DNA fragments (gBlocks Gene Fragments, IDT) containing the respective mutations, cloned into PstI and XhoI sites of the pJ4M TDP-43-TEV-MBP-His_6_-backbone.

#### Mammalian expressing constructs

To generate an expressing construct coding for Myc-hTDP-43, the coding sequence of hTDP-43 was PCR amplified from pEGFP-C1-hTDP-43 (Ederle et al., 2018), including a Myc coding sequence in the forward PCR primer, and cloned into a pcDNA5-FRT-TO-backbone using XhoI and BamHI restriction sites. Note, that the hTDP-43 template includes a resistance to TARDBPHSS118765 siRNA (Invitrogen) used to silence endogenous TDP-43 (see below “*siRNA-mediated knockdown of TDP-43*”). For generation of the TDP-43 12D and 12A constructs, synthetic gBlocks (IDT) harboring the respective mutations were previously cloned into the NdeI and BamHI sites of the pEGFP-C1-hTDP-43 vector. In constructs carrying mutations in the NLS of TDP-43 (mNLS), amino acids 82-84 as well as 95,97,98 were exchanged for alanine (pEGFP-hTDP-43 double-mNLS). Then, the mNLS region was transferred from the pEGFP-TDP-43 double-mNLS template to the pcDNA5-FRT-TO-Myc-hTDP-43, 12D and 12A vectors via the restriction enzymes XhoI and NdeI. To generate the GCR_2_-GFP_2_-TDP-43 12D and 12A constructs, the respective coding sequences were PCR amplified and inserted into GCR_2_-GFP_2_-backbone using EcoRV and BamHI. To allow for lentiviral packaging and subsequent neuronal transduction, coding sequences of TDP-43 Wt, 12D and 12A were subcloned into the FhSynW backbone in frame with mEGFP (May et al., 2014).

### Cell culture, transfection and stress treatment

HeLa cells were grown in DMEM high glucose GlutaMAX (Invitrogen) supplemented with 10% fetal bovine serum (FBS) and 10 μg/ml gentamicin and incubated in a humidified chamber with 5% CO_2_ at 37°C. cDNA transfections were performed using Lipofectamine 2000 (Thermo) in culture medium without gentamicin and medium was exchanged after 4 to 6 hours to avoid cellular stress by the transfection reagent. Note, that for equal transfection efficiency different amounts of DNA were transfected for the different constructs (For 12D: 100%; for Wt and 12A: 75% + 25% empty vector DNA). For immunostaining cells were fixed after ~24h. Hydrogen peroxide (H_2_O_2_) (1 mM) treatment was carried out for 2h, MG132 (10 μM) treatment for 2.5-3h and sodium arsenite (0.5 mM) treatment for 45 min.

### Neuronal cell culture and lentiviral packaging

Primary hippocampal and cortical neuronal cultures were prepared from embryonic day 19 rats as described in detail previously (Guo et al., 2018). In brief, neocortex and hippocampus were dissected, followed by enzymatic dissociation and gentle trituration. For immunofluorescence experiments, hippocampal neurons (85000 cells/ml) were plated on poly-D-lysine-coated glass coverslips (VWR) in 12-well plates (Thermo Fisher) and cultured in Neurobasal medium (Thermo Fisher) supplemented with 2% B27 (Thermo Fisher), 1% Penicillin-Streptomycin (Thermo Fisher), 0.5 mM L-glutamine (Thermo Fisher) and 12.5μM glutamate (Thermo Fisher). Both, cortical and hippocampal neurons, were transduced on day in vitro (DIV) 5.

Cortical neurons (250000 cells/ml) used for filter trap assays were plated on poly-D-lysine-coated 6-well plates and cultured in Neurobasal medium containing 2% B27, 1% Penicillin-Streptomycin and 0.5 mM L-glutamine.

Lentiviral packaging was performed by seeding HEK293FT cells (Thermo Fisher) of low passage number into three 10 cm dishes per construct (5×10^6^ cells/dish). Cells were plated in DMEM, high glucose, GlutaMAX (Thermo Fisher) supplemented with 10% FBS (Sigma), 1% Penicillin-Streptomycin (Thermo Fisher) and 1% Non-Essential Amino Acids (Thermo Fisher). On the following day, cells were co-transfected with 18.6 μg transfer vector (FhSynW-mEGFP-hTDP-43, FhSynW-mEGFP-hTDP-43 (12D), FhSynW-mEGFP-hTDP-43 (12A), FhSynW-mEGFP-hTDP-43-mNLS, FhSynW-mEGFP-hTDP-43-mNLS (12D), FhSynW-mEGFP-hTDP-43-mNLS (12A)), 11 μg pSPAX2 and 6.4 μg pVSVG using Lipofectamine 2000 (Thermo Fisher). The transfection media was replaced by plating media supplemented with 13 mg/mL bovine serum albumin (BSA, Sigma) on the next day. Lentivirus from the cell supernatant was collected 24h later by ultracentrifugation with a Sw28 rotor (Beckman Coulter; 22,000 rpm, 2h, 4°C). Finally, lentiviral particles were resuspended in Neurobasal media (Thermo Fisher), stored at −80°C and used for lentiviral transduction by adding to neuronal culture media upon thawing. Neurons were kept in culture for 4 additional days after transduction on DIV5 (DIV5+4).

### Recombinant protein expression and purification

#### TDP-43-TEV-MBP-His_6_

All TDP-43-MBP-His_6_ variants were purified according to (Wang et al., 2018) with minor adaptations. First, expression of proteins was performed in *E. coli* BL21-DE3 Rosetta 2 using 0.5 mM IPTG and grown overnight at 16°C. Next, cells were resuspended in lysis buffer (20 mM Tris pH 8, 1 M NaCl, 10 mM imidazole, 10 % (v/v) glycerol, 4 mM β-mercaptoethanol and 1 μg/ml each of aprotinin, leupeptin hemisulfate and pepstatin A) supplemented with 0.1 mg/ml RNase A, and lysed using lysozyme and sonication. Subsequently, the protein was purified by Ni-NTA agarose (Qiagen) and eluted with lysis buffer containing 300 mM imidazole. For all TDP-43-MBP-His_6_ variants a final size exclusion chromatography (SEC) (Hiload 16/600 Superdex 200 pg, GE Healthcare) purification step was performed in purification buffer (20 mM Tris pH 8, 300 mM NaCl, 10% (v/v) glycerol supplemented with 2 mM TCEP), in order to separate TDP-43-MBP-His_6_ from protein aggregates and contaminants. Purified monomeric TDP-43-MBP-His_6_ was collected by pooling the fractions corresponding to peak B in the SEC profile (figure S1D). All purified proteins were concentrated using Amicon ultra centrifugal filters and then flash frozen and stored at −80°C. To determine protein concentration, absorbance at 280 nm was measured using the respective extinction coefficient (ε) predicted by the ProtParam tool. Additionally, for all purified proteins, the A260/280 ratio was determined and found to be between 0.5-0.7.

#### CK1δ

The kinase domain of CSNK1D was expressed as an N-terminal MBP-tagged fusion in E. coli Rosetta 2 cells, co-expressing λ-phosphatase to guarantee a completely unphosphorylated protein. The cells were grown to an OD of 0.45 and subsequently the temperature was reduced to 18°C. Then the cells were induced (generally at OD 0.7-0.8) with 0.5 mM IPTG and expression was performed overnight. Cells were harvested and resuspended in AC-A buffer (25 mM Bis-Tris, 500 mM NaCl, 10 mM β-mercaptoethanol, pH 7.0), supplemented with DNAse, RNAse, lysozyme and protease inhibitor cocktail (selfmade) for cell disruption. Lysis was done by sonication on ice (5×30 s with breaks of 1 min between each pulse). Cell debris was pelleted by centrifugation (SS34 rotor, 17 000 r.p.m., 30 min). The supernatant was filtered and subsequently loaded on a Dextrin Sepharose column (cytiva), previously equilibrated with AC-A buffer. The column was washed for 5 column volumes with AC-A buffer. Elution was done with MBP-B buffer (25 mM Bis-Tris, 500 mM NaCl, 10 mM β-mercaptoethanol, 20 mM Maltose, pH 7.0). The eluted protein was subject to TEV protease cleavage overnight at 4°C. On the next day the buffer was exchanged to IEX-A buffer (25 mM Bis-Tris, 50 mM NaCl, 10 mM. β-mercaptoethanol) by ultra-filtration (Amicon Ultra-15 30 kDa, Merck Millipore) and subject to cation-exchange chromatography by a linear to IEX-B buffer (25 mM Bis-Tris, 500 mM NaCl, 10 mM. β-mercaptoethanol). Eluted protein was concentrated and gel-filtered over a Superdex 75 (cytiva) into SEC buffer (25 mM Bis-Tris, 50 mM NaCl, 10 mM MgCl2, 1 mM DTT). Fractions were collected, concentrated and aliquots of 200 μl were flash frozen and stored at −80°C until use.

#### His_6_-TEV protease

His_6_-TEV protease expression and purification was performed as described in (Hutten et al., 2020).

### *In vitro* phosphorylation

TDP-43-MBP-His_6_ was *in vitro* phosphorylated with CK1δ and 200 μM Adenosine triphosphate (ATP) in phosphorylation buffer (50 mM Tris-HCl, pH 7.5, 10 mM MgCl2, 1 mM DTT) for 2 h at RT, using a 2-fold molar excess of TDP-43-MBP-His_6_ over CK1δ. Subsequently, the reaction was used for sedimentation and SG association assays. As negative controls, either the kinase or the ATP was omitted and also included as controls in subsequent assays.

### Enzymatic digestion, enrichment for phosphopeptides and mass spectrometric analysis

TDP-43-MBP-His_6_ was *in vitro* phosphorylated as described above, separated on a 10 % SDS-PAGE gel and visualized by Coomassie staining. The gel band corresponding to the phosphorylated TDP-43-MBP-His_6_ was excised and destained twice for 30 min at 37 °C with 50 % acetonitrile in 50 mM Tris-HCl, pH 8. The gel piece was dehydrated with 100 % acetonitrile, reduced and alkylated, and finally digested overnight at 37 °C with 375 ng Trypsin (Promega). The peptides were extracted from the gel twice using 100 μl of 50 % acetonitrile and 0.25 % TFA buffer. Both extractions were merged and evaporated in a vacuum evaporator. In order to enrich the phosphopeptides, 10 μl of 0.5 mg/μl TiO2 beads (GL Sciences Cat.No: 5010-21315) in loading buffer (80% ACN, 5% TFA, 1M Glycolic acid) were added to the dried samples in a ratio of 0.3 mg of beads to 5 pmol of protein. Samples were incubated for 10 min at RT on a shaker at 2000 rpm and spun down at 1200 rpm for 1 min. The supernatant was removed and kept for further analysis, while beads were sequentially washed with loading buffer, washing buffer 1 (80% ACN, 1% TFA) and washing buffer 2 (10% ACN, 0.2% TFA). Next, the beads were dried in the hood for 10 min and resuspended with 50 μl elution buffer (28 % ammonia solution in H_2_O). Finally, the samples were speed vacuum evaporated and resuspended with 15 μl 0,1 % FA. For LC-MS purposes, desalted peptides were injected in an Ultimate 3000 RSLCnano system (Thermo) and separated in a 25-cm analytical column (75μm ID, 1.6μm C18, IonOpticks) with a 30-min gradient from 3 to 30% acetonitrile in 0.1% formic acid. The effluent from the HPLC was directly electrosprayed into a Qexactive HF (Thermo) operated in data dependent mode to automatically switch between full scan MS and MS/MS acquisition. Survey full scan MS spectra (from m/z 300-1600) were acquired with resolution R=60,000 at m/z 400 (AGC target of 3×10_6_). The 10 most intense peptide ions with charge states between 2 and 5 were sequentially isolated to a target value of 1×10^5^ with resolution R=15,000 and isolation window 1.6 Th and fragmented at 27% normalized collision energy. Typical mass spectrometric conditions were: spray voltage, 1.5 kV; no sheath and auxiliary gas flow; heated capillary temperature, 250°C; ion selection threshold, 33.000 counts.

### Fluorescent labeling of purified TDP-43

TDP-43-MBP-His_6_ variants were labeled with Alexa Fluor 488 C5 maleimide (Thermo Fisher) at a low (~0.01-0.05) labelling efficiency in order to avoid interference with condensate formation. Labeling was performed according to the manufacture’s protocol using a 1:100 or 1:20 protein:fluorescent dye mole ratio. Briefly, the Alexa Fluor reagent, previously dissolved in DMSO, was mixed with the protein and kept in the dark for 2 h at RT. Excess dye was removed by consecutive washes with TDP-43 purification buffer using Amicon ultra centrifugal filters. Subsequently, labelled protein was used for spinning disc confocal microscopy, FRAP and aggregation assays, respectively.

### *In vitro* phase separation and aggregation assays

#### Sedimentation assay

For sedimentation analysis, 1 μM TDP-43-TEV-MBP-His_6_ variants or *in vitro* phosphorylated TDP-43-TEV-MBP-His_6_ was cleaved by addition of 20 μg/ml His_6_-TEV protease in 50 μl or 25 μl Hepes buffer (20 mM Hepes, pH 7.5, 150 mM NaCl, 1 mM DTT), respectively, to remove the MBP-His_6_ tag and induce phase separation. Samples were incubated for 60 min at 30 °C, followed by centrifugation for 15 min at 21,000 g at 4°C to pellet the formed condensates. Equal amounts of supernatant (S) and condensate (C) fractions were loaded onto an SDS-PAGE gel and TDP-43 was detected by Western Blot (rabbit TDP-43 N-term, Proteintech, Cat.No: 10782-2-AP).

#### Microscopic condensate assay

For all microscopic condensate assays Uncoated μ-Slide 18 Well - Flat chambers (Cat.No: 81821, Ibidi) were pretreated with 10% Pluronics F-127 solution for 1h and 5 times washed with MilliQ water. The water remained in the chamber until just before the experiment, as described in (Ceballos et al., 2018).

Purified TDP-43-TEV-MBP-His_6_ variants were buffer exchange to Hepes buffer or phosphate buffer (20 mM Na_2_HPO_4_/NaH_2_PO_4_, pH 7.5, 150 mM NaCl, 2.5% glycerol, 1 mM DTT). Proteins were then centrifuged at 21,000 g for 10 min at 4°C to remove any preformed protein precipitates. For condensates formation, the reaction was setup directly in Pluronics-coated μ-Slide 18 Well - Flat chambers, where proteins were diluted to the indicated concentrations and phase separation was induced by addition of 100 μg/ml His_6_-TEV protease at RT. After ~20 min, imaging was performed by bright field microscopy using a widefield microscope.

For fusion events and FRAP analysis, condensates were formed directly in Pluronics-coated μ-Slide 18 Well - Flat chambers as described above using 20 μM of each Al.488-labelled TDP-43 protein variants (Wt, 5D, 12D) in Hepes buffer and incubated for 10 min at RT before imaging. Note that experiments were performed until maximally 1 h after adding the TEV protease, in order to avoid *in vitro* aging of condensates.

#### Turbidity assay

Phase separation of TDP-43-TEV-MBP-His_6_ variants was induced as described above for the microscopic condensate assay. Reactions of 20 μl samples were prepared at the indicated concentrations in 384-well plates and incubated for 30 min at RT after adding TEV protease. Subsequently, a BioTek Power Wave HT plate reader was used to measure turbidity at 600 nm. Turbidity measurements were performed in triplicates.

#### Semi-denaturing Detergent Agarose Gel Electrophoresis (SDD-AGE)

SDD-AGE experiments were performed based on protocols published by (French et al., 2019) and (Halfmann and Lindquist, 2008). First, 2 μM purified TDP-43-MBP-His_6_ variants (WT, 5D, 12D, 12A) were set up in low binding tubes (Eppendorf) in 35 μl aggregation buffer (50 mM Tris pH 8.0, 250 mM NaCl, 5% glycerol, 5% sucrose, 150 mM imidazole pH 8.0) and supplemented with 1x protease inhibitor (Sigma). Samples were shaken for 30 min at 1000 rpm at RT (~22°C) and then incubated at RT for the indicated time period. 5 μl of each sample was collected and diluted in SDD-AGE buffer (40 mM Tris-HCl pH 6.8, 5% Glycerol, 0.5% SDS, 0.1% bromphenol-blue) and analyzed by SDD-AGE by horizontal 1.5% agarose gel electrophoresis (gel: 1.5% agarose in 20 mM Tris, 200 mM Glycine and 0.1% SDS) in running buffer (60 mM Tris, 20 mM Acetate, 200 mM glycine, 1 mM EDTA and 0.1% SDS) for ~6 h at 60 V. Detection of TDP-43 monomers, oligomers and high molecular weight species was performed after overnight capillary transfer in TBS (50 mM Tris pH 7.6, 150 mM NaCl) to a nitrocellulose membrane and by standard Western Blot using rabbit anti TDP-43 N-term antibody (Proteintech, Cat.No: 10782-2-AP).

### Formation of Alexa 488-labelled TDP-43 aggregates

In order to visualize TDP-43 (wt, 5D, 12D and 12A) aggregates formed under the above described assay conditions, 10 μM Al.488-labeled TDP-43-MBP-His_6_ was set up in low binding tubes (Eppendorf) in aggregation buffer and incubated with or without 100 μg/ml His_6_-TEV protease. Samples were shaken at 1000 rpm at RT for 30 min and then transferred into a 384-well black plate (Greiner Bio-One), incubated at RT and imaged by confocal microscopy after 2, 8 and 24h.

### Cellular TDP-43 solubility assays

#### Fractionation in RIPA-Benzonase buffer

HeLa cells (~1×10^6^) were washed twice in PBS, harvested by scraping and pelleted at 3600 rpm for 5 min. Cell pellets were incubated on ice for 15 min in 200 μl RIPA buffer (50 mM Tris-HCl, pH 8.0, 150 mM NaCl, 1% NP-40, 0.5% deoxycholate, 0.1% SDS) with freshly added 1x protease inhibitor cocktail (Sigma), 1x phosphatase inhibitors (final concentration: 10 mM NaF, 1 mM β-glycerophosphate, 1 mM Na3VO4) and 0.05 unit/μl Benzonase (Sigma). Samples were sonicated in a BioRuptorPico (Diagenode) for 45 sec and 20 μl of sample was collected as “Input”. The remaining sample was then centrifuged at 13,000 g for 30 min at 4°C. The resulting supernatants (S) were collected and the remaining pellets were washed in RIPA buffer with inhibitors, resonicated for 45 sec and recentrifuged for 30 min at 4°C at 13,000 g. Finally, the RIPA insoluble pellets (I) were resuspended in 36 μl Urea buffer (7 M urea, 2 M thiourea, 4% CHAPS, 30 mM Tris-HCl, pH 8.5) and sonicated. All samples were supplemented with 4x Lämmli buffer (62,5 mM Tris HCl, pH 6,8, 10% glycerol, 2% SDS, 0,03% bromphenol-blue, 143 mM β-Mercaptoethanol) and Input and Supernatant (S) samples were boiled prior to SDS-PAGE and Western Blot against TDP-43 (rabbit anti TDP-43 C-term, Proteintech, Cat. No:12892-1-AP) and Myc (mouse anti-myc 9E10 antibody, Helmholtz Center Munich). Note that for detection reasons, the (I) fractions were 4x more concentrated than the (S) fractions, so they are represented in a 1:5 ratio.

#### Filter trap assay

Cortical neurons expressing the indicated GFP-tagged TDP-43 variants (DIV5 + 4 days expression) were washed two times with PBS and lysed on ice in RIPA buffer (50 mM Tris-HCl, pH 8.0, 150 mM NaCl, 1% NP-40, 0.5% deoxycholate, 0.1% SDS) freshly supplemented with 1x protease inhibitor cocktail (Sigma), 1x phosphatase inhibitor cocktail (Sigma) and 0.125 Units/μL Benzonase (Sigma) for 20 min. Cell lysates were collected and centrifuged at 1,000 g, 4°C for 30 min. 2/3 of the resulting supernatant (RIPA-insoluble fraction) was filtered through a nitrocellulose membrane (0.2 μM pore size, Merck) using a filter trap slot blot (Hoefer Scientific Instruments). After washing with PBS for three times, membranes were blocked for 1h with 2% I-Block (Thermo Fisher) prior to immuno-detection with mouse anti-GFP (UC Davis/NIH Neuromab Facility, Cat.No: N86/8) and rabbit anti Calnexin antibody (Enzo Life Sciences, Cat.No: ADI-SPA-860). The remaining 1/3 of the lysates was diluted with 3x loading buffer (200 mM Tris-HCl pH 6.8, 6% SDS, 20% glycerol, 0.1 g/ml DTT, 0.1 mg Bromophenol Blue), boiled at 95°C and used for subsequent standard Western Blot analysis.

### Multi scale simulations

#### Coarse-grained MD simulations

Coarse-grained MD simulations with explicit solvent to investigate protein phase separation and phase-separated protein condensates were run with a rescaled version of the Martini forcefield (Marrink et al., 2007; Monticelli et al., 2008) as described by (Benayad et al., 2021). A similar approach was shown to describe the conformational ensembles of proteins with disordered domains very well (Larsen et al., 2020; Martin et al., 2021), and we recently showed that such approaches can be extended to simulations of liquid-liquid phase separation of disordered proteins (Benayad et al., 2021). Protein-protein interactions were thus scaled to 0.8 of the default value. Chloride and sodium ions were added to neutralize the system in simulations of Wt and 12D proteins. 10% of the water beads were replaced by WF anti-freeze beads. Coarse-grained simulations were run with GROMACS 2018 (Abraham et al., 2015). Simulations boxes measured 450 x 450 x 600 Å. Simulations systems were energy minimized and equilibrated in MD simulations with and without position restraints. 118 Wt and 12D C-terminal LCDs (aa. 261 – 414) were simulated for 20 μs each. The coarse-grained simulations systems consist of roughly one million particles. Equations of motions were integrated with a 20 fs time step. Simulations were conducted in the NPT ensemble at 1 bar and 300 K using the Parrinello-Rahman barostat (Parrinello and Rahman, 1981) and the Bussi-Donadio-Parrinello velocity-rescaling thermostat (Bussi et al., 2007).

Note that in the coarse-grained approach we employed, four atoms are typically group together to a coarse-grained particle. E.g., a coarse-grained water molecule would correspond to four water molecules in an atomistic simulation.

#### Atomistic MD simulations

Hierarchical chain growth (HCG) (Pietrek et al., 2020) enables us to generate statistically independent and chemically-meaningful conformations of a biomolecular condensate with atomic resolution, which serve as starting points for atomistic MD simulations. Atomic-resolution models of clusters of the C-terminal disordered domain of TDP-43 (aa. 261 – 414) were generated for both Wt protein and the 12D mutant. To assemble disordered proteins into a condensate, we first assemble pairs of disordered domains, then pairs of pairs, pairs of quadruplets, and so forth, following the logic set out in (Pietrek et al., 2020). HCG Monte Carlo manifestly satisfies detailed balance and thus we generate representative ensembles. Finally, we arrive at densely packed disordered domains, while retaining atomic resolution at each modeling step. Periodic boundary conditions were employed during the assembly.

Clusters of Wt and 12D LCDs were solvated in a 150 Å x 150 Å x 150 Å simulation box, the system charge was neutralized and 150 mM NaCl was added. We employed the Amber-disp protein force field developed by Robustelli and et al. (Robustelli et al., 2018), including the modified TPIP4P-D water model (Piana et al., 2015) that accompanies the Amber-disp protein force field. Temperature was maintained at 300 K by the Bussi-Donadio-Parrinello velocity-rescaling thermostat (Bussi et al., 2007). We employed the Parrinello-Rahman barostat (Robustelli et al., 2018) to set the pressure to 1 bar. Equations of motions were integrated with a 2 fs time step. Production simulations were prepared by energy minimization with and without soft-core potentials. To start production simulations, we equilibrated the atomistic simulations systems, running at least 5000 steps with a 1 fs time step and position restraints and for 1.5 ns with a 2 fs time step also with position restraints. Equilibrium simulations of the clusters of the disordered domains were conducted with GROMACS 2019 (Abraham et al., 2015). For both wild-type and 12D clusters of 32 chains with 154 residues were simulated for 1 μs with two repeats each started from independently generated HCG structures.

Simulations were analyzed with the MDAnalysis (Gowers et al., 2016; Michaud-Agrawal et al., 2011) and the MDtraj Python libraries (McGibbon et al., 2015). Contact analysis was performed with the Contact Map Explorer Python library (https://github.com/dwhswenson/contact_map).

### Nuclear transport assay

To analyze import of GCR_2_-GFP_2_ tagged TDP-43 reporters, HeLa cells were grown for at least 2 passages in DMEM supplemented with 10% dialyzed FBS and were transiently transfected with the different GCR_2_-GFP_2_-TDP-43 variants as described above. Import of the GCR_2_-GFP_2_-TDP-43 reporters was induced by adding dexamethasone (5 μM final concentration) in imaging medium (fluorobrite) and followed by live cell imaging using a spinning disk confocal microscope.

### SG association assay

HeLa cells were grown on high precision (No. 1.5) poly-L-lysine coated 12 mm coverslips, and after SG induction with MG132 (10 μM for 2.5-3h), cells were permeabilized for 2x 2 min with 0.004-0.005% digitonin in KPB (20 mM potassium phosphate pH 7.4, 5 mM Mg(OAc)_2_, 200 mM KOAc, 1 mM EGTA, 2 mM DTT and 1 mg/ml each aprotinin, pepstatin and leupeptin). Soluble proteins were removed by several, stringent washes (4x 4 min in KPB on ice) before blocking nuclear pores by 15 min incubation with 200 mg/ml wheat germ agglutinin (WGA) on ice. Cells were then incubated for 30 min at RT with 100 nM TDP-43-MBP-His_6_. For comparison of *in vitro* phosphorylated TDP-43 with controls, proteins were either subjected to the *in vitro* phosphorylation reaction or mock treated (Wt, 12D) in absence of kinase or ATP before exchanging the buffer to KPB using 40K Zeba spin desalting columns (Thermo). Subsequently, cells were washed (3x 5 min in KPB on ice) to remove unbound TDP-43-MBP-His_6_ and processed by immunostaining to visualize SGs. SGs and TDP-43-MBP-His_6_ were visualized by G3BP1 immunostaining (rabbit anti G3BP1antibody, Proteintech, Cat.No: 13057-2-AP) and MBP immunostaining (by mouse anti MBP antibody, Proteintech, Cat.No: 66003-1-Ig), respectively. On Fig. 4B for clarity, signals were converted to grey values in the individual channels (upper two rows). In the merge (lower row), G3BP1 is shown in magenta, TDP-43-MBP-His_6_ in green, white pixels indicate colocalization. Nuclei were counterstained with DAPI (turquoise).

### siRNA-mediated knockdown of TDP-43

TDP-43 knockdown was achieved using the pre-designed TARDBPHSS118765 siRNA (Invitrogen) as described in (Dormann et al., 2009). Briefly, 20 nM siRNA was transfected into HeLa cells using RNAimax (Thermo) transfection reagent. Knockdown was analyzed 48 h post transfection by immunohistochemistry using mouse anti TDP-43 antibody (Proteintech, Cat.No: 60019-2-Ig) and immunoblotting using rabbit anti TDP-43 C-Term antibody (Proteintech, Cat.No: 12892-1-AP) to detect TDP-43 and mouse anti alpha-Tubulin antibody (Proteintech, Cat.No: 66031-1-Ig) for detection of α-Tubulin as a control.

### RNA extraction and RT-PCR to analyze TDP-43 splice targets

TDP-43 expression was silenced in HeLa cells by siRNA as described above and 24 h later cells were transfected with siRNA-resistant pcDNA5-FRT-TO-Myc-hTDP43 constructs (Wt, 12D and 12A). 48h after transfection, cells were harvested and total RNA was extracted using an RNeasy mini kit from Qiagen. cDNA was synthesized using 500 ng of total RNA, M-MLV reverse transcriptase polymerase (Invitrogen), and 6 μM of random hexamer primer (NEB). cDNA was amplified with Taq DNA polymerase (NEB) using the forward (FW) and reverse (RV) primers targeting the *SKAR* gene (FW - 5’CCTTCATAAACCCACCCATTGGGACAG3’; RV-5’GTGGTGGAGAAAGCCGCCTGAG3’) (Fiesel et al., 2012) and the *BIM* gene (FW-5’TCTGAGTGTGACCGAGAAGG3’; RV - 5’TCTTGGGCGATCCATATCTC 3’) (Tollervey et al., 2011). PCR products were separated by electrophoresis on a 2.5% agarose gel containing GelRed (Sigma).

### Electrophoretic-mobility shift assays (EMSA)

The TDP-43 autoregulatory RNA site (Ayala et al., 2011) located in the *TARDBP* 3’UTR (5’UCACAGGCCGCGUCUUUGACGGUGGGUGUCCCAUUUUUAUCCGCUACUCUUUAUUUCAUGGAGUCGUAUCAACGCUAUGAACGCAAGGCUGUGAUAUGGAACCAGAAGGCUGUCUGAACUUUUGAAACCUUGUGUGGGAUUGAUGGUGGUGCCGAGGCAUGAAAGGCUAGUAUGAGCGAGAAAAGGAGAGAGCGCGUGCAGAGACUUGGUGGUGCAUAAUGGAUAUUUUUUAACUUGGCGAGAUGUGUCUCUCAAUCCUGUGGCUUUGGUGAGAGAGUGUGCAGAGAGCAAUGAUAGCAAAUAAUGUACGAAUGUUUUUUGCAUUCAAAGGACAUCCACAUCUGUUGGAAGACUUUUAAGUGAGUUUUUGUUCUUAGAUAACCCACAUUAGAUGAAUGUGUUAAGUGAAAUGAUACUUGUACUCCCCCUACCCCUUUGUCAACUGCUGUG) was *in vitro* transcribed from double-stranded DNA templates and Cy5-labeled using the HyperScribe™ T7 High Yield Cy5 RNA Labeling Kit (APExBIO, Cat.No: K1062) per manufacturer’s instructions. (UG)_12_ RNA (5’ UGUGUGUGUGUGUGUGUGUGUGUG) was chemically synthesized with the addition of a 5’ Cy5-label (Metabion). 16 nM of Cy5-labeled RNA was mixed with varying amounts of TDP-43 Wt, 12A, and 12D (0-1.6 μM). Binding reactions (20 μL) were incubated in binding buffer [20 mM NaPO_4_ (pH 8), 150 mM NaCl, 1 mM DTT, 5 mM MgCl_2_, 0.5 mg/mL BSA, 0.1 mg/mL yeast tRNA, 5% glycerol, 1 U/μL RNase Inhibitor (NEB)] for 20 minutes at RT before loading onto a 1 mm thick non-denaturing polyacrylamide gel (6%) in 0.5x TBE. Gels were run at 100 V for 1 hour at RT. Gels were imaged with a Typhoon™ FLA 9500 laser scanner.

### Immunostaining

All steps were performed at RT. HeLa cells were fixed in 4% formaldehyde in PBS for 10 min, permeabilized for 5 min using 0.2% (v/v) Triton X-100 in PBS and subsequently blocked in blocking buffer (5% goat or donkey serum in 0.1% saponine in PBS) for 30 min. Primary and secondary antibodies were diluted in blocking buffer and applied each for 1h and washed three times using 0.1% saponine in PBS. TDP-43 was stained using mouse anti TDP-43 antibody (Proteintech), SGs were stained using goat anti TIA1 antibody (Santa Cruz, Cat. No: sc-48371) or rabbit anti G3BP1 antibody (Proteintech) and DNA was stained with DAPI at 0.5 μg/ml in PBS for 5 min. Coverslips were then mounted on glass slides with ProLong™ Diamond Antifade reagent (Life Technologies) and dried overnight at RT.

Hippocampal neurons cultured on glass coverslips were washed twice with PBS and fixed for 10 min at RT using 4% paraformaldehyde and 4% sucrose in PBS. Primary antibody as well as secondary antibody (1:400) were diluted in GDB buffer (0.1% gelatin, 0.3% Triton X-100, 450 mM NaCl, 16 mM sodium phosphate pH 7.4). Primary antibody (Mouse anti Map2, Sigma, Cat# M1406, RRID: AB_477171) was incubated overnight at 4°C while secondary antibodies was applied for 1h at RT, each followed by three times washing with PBS. Coverslips were mounted using Vectashield Vibrance with DAPI (Biozol) to counterstain nuclei.

### Microscopy

#### Bright and wide-field microscopy

Imaging of unlabeled TDP-43 condensates was done by bright-field microscopy on an Axio Oberver.Z1 wide-field fluorescence microscope, using a 63x/1.40 Oil objective and an AxioCam 506 (Zeiss, Oberkochen, Germany).

#### Confocal microscopy

Confocal microscopy was performed using an inverted Leica SP8 microscope and the LAS X imaging program (Leica), provided by the Bioimaging core facility of the Biomedical Center, which included the excitation lasers for 405, 488, 552 and 638 nm. Images were acquired using two-fold frame averaging with a 63×1.4 oil objective, with an image pixel size of 180 nm for Al.488-TDP-43 aggregates and fixed cells, and 59 nm for images of cells subjected to the SG association assay. The following fluorescence settings were used for detection: DAPI: 419-442 nm, GFP: 498-533 nm, Alexa 555: 562-598 nm, Alexa 647: 650-700 nm. Recording was performed sequentially using a conventional photomultiplier tube (PMT) to avoid bleed-through.

#### Spinning disc confocal microscopy

##### a) Nuclear transport assay imaging

Images were acquired for a duration of 50 min in 2.5 min intervals at 36.5°C and 5% CO_2_ (EMBLEM environmental chamber) using an inverted microscope (Axio Observer.Z1; Carl Zeiss, Oberkochen, Germany) equipped with a confocal spinning disc (CSU-X1; Yokogawa, Tokyo, Japan) and a 63x/1.4 oil immersion lens. Images were acquired using the 488 nm SD laser line and an EM-CCD camera (EvolveDelta; Photomoetrics) at bin 1×1.

##### b) Fusion events and Fluorescence Recovery after Photobleaching (FRAP)

Experiments were performed on an inverted microscope (Axio Observer.Z1; Carl Zeiss, Oberkochen, Germany) equipped with a confocal spinning disk unit (CSU-X1; Yokogawa, Tokyo, Japan) and an oil immersion lens of 100x/1.46 Oil Ph3. Images recording the dynamics of TDP-43 condensates were obtained using a EM-CCD camera (EvolveDelta; Photomoetrics), with a bin 1×1 in a recording mode of 5 sec intervals in a block of 3 min. Images of TDP-43 condensates after bleaching were acquired with bin 1×1 in streaming mode for 1.5 sec followed by a block of 2 min where images were recorded in intervals of 5 sec. Experiments were performed at RT and ≥11 condensates were analyzed per condition in three independent experiments. Localized photobleaching (“half-bleach”) was obtained using a laser scanning device (UGA-42 Geo; Rapp OptoElectronic, Hamburg, Germany). The “Geo” module allowed for simultaneous laser illumination within hardware-defined shapes of different sizes. For this experiment, an illumination size of ~4 μm in a square-like shape was used. The targeting structure was half bleached to approximately 70% of the initial intensity using a 473 nm diode laser (DL-473/75; Rapp OptoElectronic, Hamburg, Germany).

### Quantification and analysis

#### Droplet quantification

Wide-field images of droplets were processed and quantified and measured using Image J/Fiji software. First, a bandpass filter of 20 pixels was applied to all images in order to reveal some details and thresholds were adjusted to optimally include all droplets. Finally, droplets were counted and measured by their size and roundness [4*area/(π*major_axis^2), or the inverse of the aspect ratio] using the command Analyze Particles, excluding the detection of particles with a circularity below 0.3 and/or an area smaller than 3 pixels. Statistical analyses were performed in GraphPad Prism 8.

#### Analysis of cellular images

Analysis of the nuclear transport assay was performed using Image J/Fiji software by measuring loss in cytoplasmic fluorescence over time and normalizing t= 0 min to 1.

Images of cells from the SG association assay (Hutten and Dormann, 2020) were processed and analyzed using Image J/Fiji software, applying linear enhancement for brightness and contrast and implemented plugins for measurement of pixel intensities in SGs.

Quantification of Myc-hTDP-43 recruitment into SGs was performed using Image J/Fiji software. First, SGs from TDP-43-positive cells were selected using the Wand tracing tool and a band of 1 μm representing a proxy for the cytosol was drawn around all selected SGs using the ‘Make Band’ command. Then, all pixel intensities for both SG and band selections was extracted for the TDP-43 channel. After subtraction of the background signal from all measured values, calculation of the SG/band ratio was performed for each SG and normalized SG/band ratio = 0 to 1.

Analysis of Myc-hTDP-43 recruitment into NBs was performed by counting the number of cells with positive TDP-43 nuclear condensates, excluding cells expressing TDP-43 staining only in the cytoplasm. Profile of TDP-43 nuclear staining was performed using Image J/Fiji software by using the ‘Plot Profile’ command, which quantifies the gray values along the indicated lines.

All statistical analyses were performed in GraphPad Prism 8.

#### FRAP analysis

FRAP analysis were performed using Image J/Fiji software by calculating the fluorescence intensity over time (I(t)) using the macro Time Series Analyzer command and the following formula:

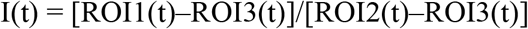

ROI1 corresponds to the averaged gray values of the bleached area, and ROI2 to the averaged gray values of the total droplet. ROI3 corresponds to the averaged background values. Values were further normalized to the initial fluorescence by dividing I(t) by the mean gray value of the initial 1 time step before bleaching <I(1)>. This way bleached areas were corrected for background noise and bleaching due to imaging over time.

---

**Fig. S1.**
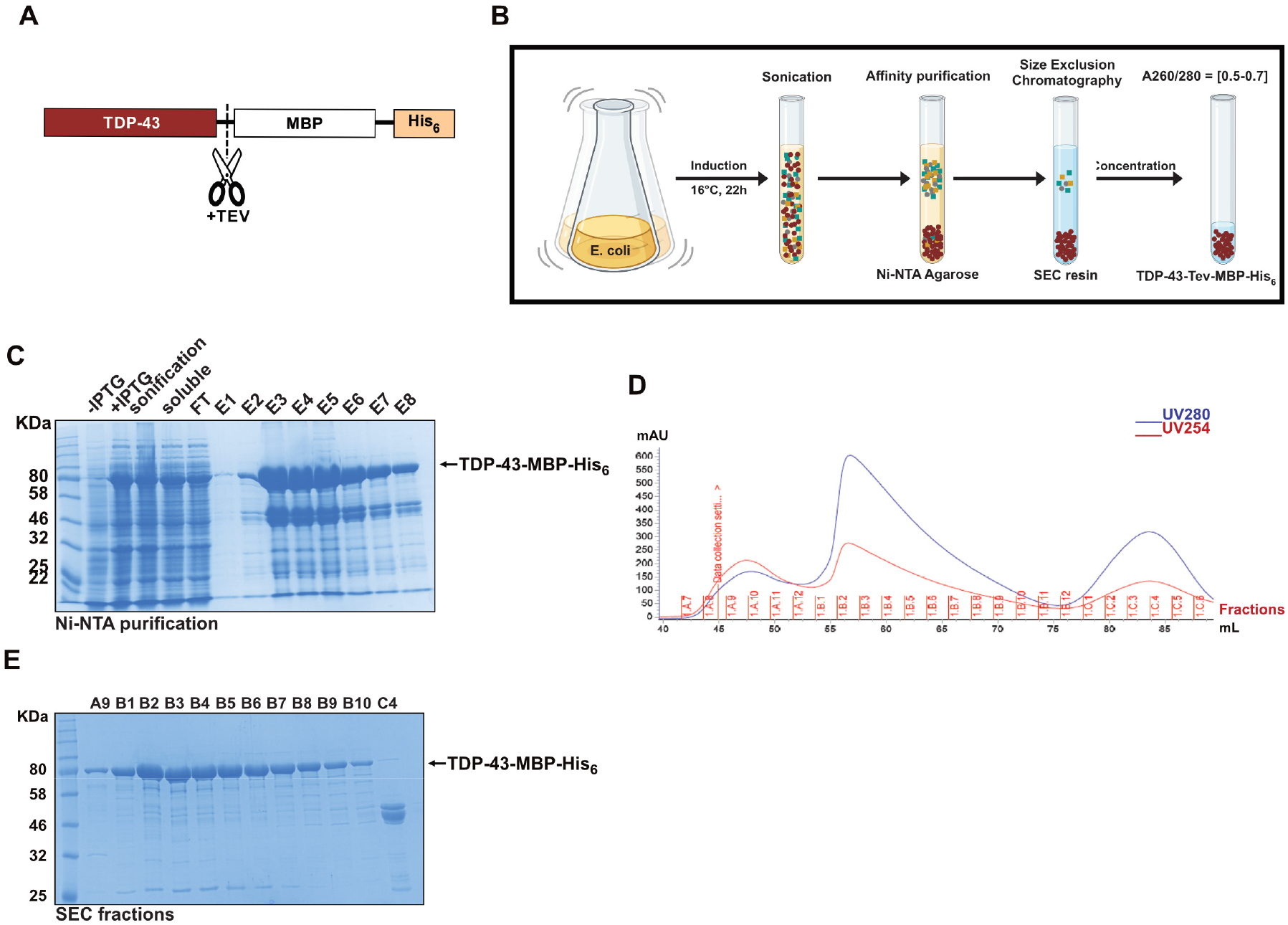
TDP-43-MBP-His_6_ purification. A Schematic representation of purified full-length TDP-43 containing a solubilizing maltose binding protein (MBP) tag, a His_6_-tag and a TEV protease recognition site before the MBP tag. B Schematic diagram of TDP-43-MBP-His_6_ expression and purification (created in BioRender.com). First, the construct is expressed in *E. coli* for 22 h at 16 °C and bacteria are lysed by sonication. Finally, the protein is purified under high salt conditions via Ni-NTA affinity purification and size exclusion chromatography (SEC) to obtain clean soluble TDP-43-MBP-His_6_ (red circles), which is largely devoid of nucleic acids, as judged from the A260/280 ratio. C Representative SDS-PAGE gel of different steps from protein expression to Ni-NTA affinity purification. First and second lines correspond to samples before (-IPTG) and after (+IPTG) induction of TDP-43-MBP-His_6_ expression. Third, fourth and fifth line correspond to samples after sonication, supernatant collection after spin down (soluble) and flow through (FT) after first wash with lysis buffer. The last 8 lines correspond to the consecutive elution steps (E1-E8) from the Ni-NTA agarose beads, which were pooled (E2-E6) and used for SEC. ~89 kDa bands were detected for TDP-43-MBP-His_6_ using Coomassie staining. D Representative SEC profile showing the amount of protein (UV280) and nucleic acids (UV254) on the y-axis versus the volume (mL) of the different fractions after SEC on the x-axis. E Coomassie-stained SDS-PAGE gel showing the indicated fractions (A9-C4) of the SEC run displayed in (D). A9, B1-B10 and C4 are fraction samples from peaks A, B and C in the SEC profile (D) that represent TDP-43-MBP-His_6_ oligomeric, monomeric and cleaved species, respectively.

**Fig. S2.**
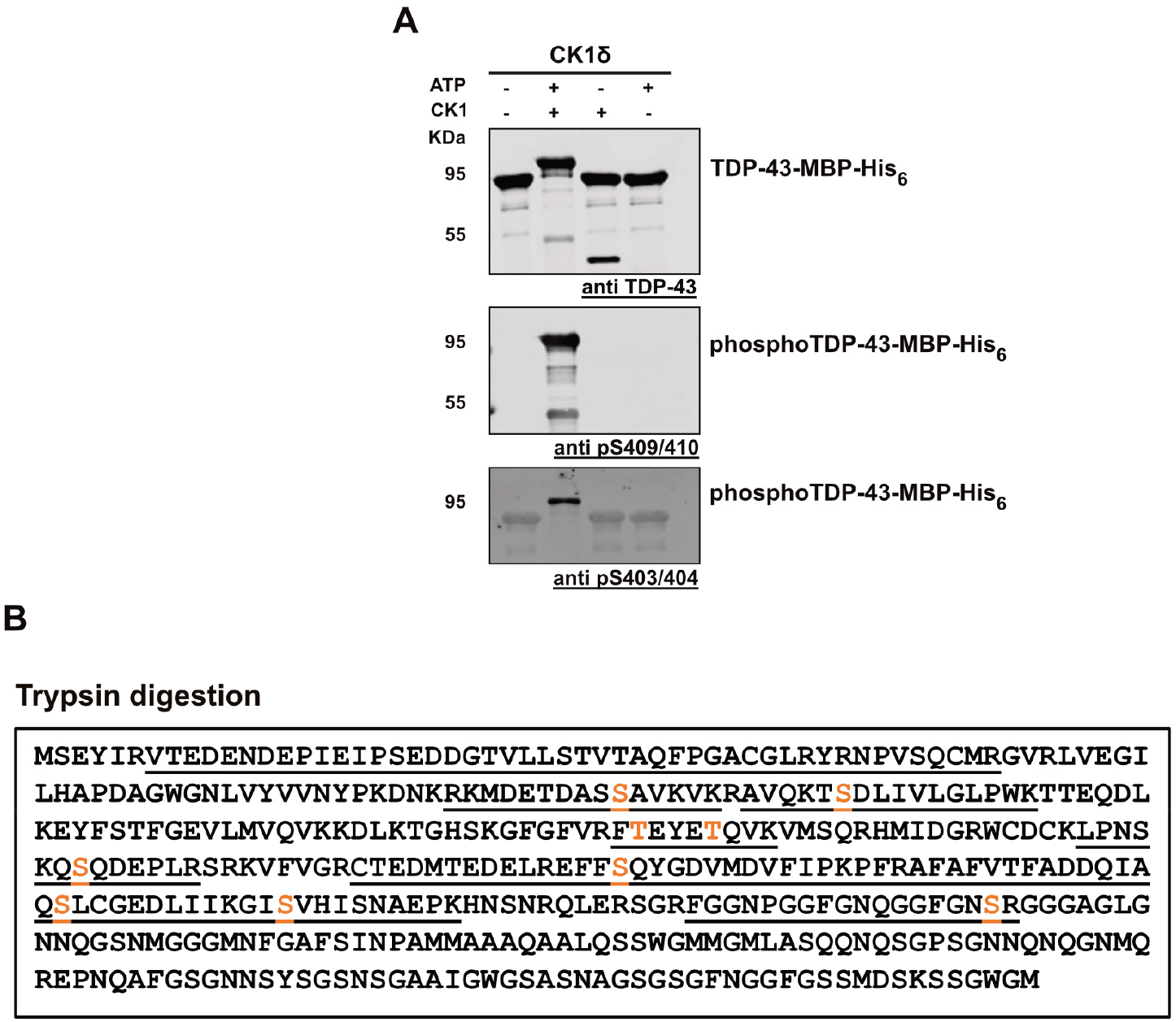
Identification of TDP-43-MBP-His_6_ phospho-sites after *in vitro* phosphorylation with CK1δ. A Identification of TDP-43 phospho-sites on *in vitro* phosphorylated TDP-43 (+CK1δ, +ATP) in comparison to controls (-CK1δ-ATP; CK1δ only; ATP only) by Western blot. Samples were run on SDS-PAGE and analyzed by Western blot using a rabbit anti-TDP-43 N-term antibody (Proteintech) to detect total TDP-43, rat anti-TDP-43-phospho Ser409/410 (clone 1D3, Helmholtz Center Munich) and mouse anti-TDP-43-phospho Ser403/404 (Proteintech, Cat.No: 66079-1-Ig) antibodies. B Schematic diagrams showing sequence coverage in mass spectrometry after trypsin digest (underlined) and phosphorylated serine/threonine residues (orange) of *in vitro* phosphorylated TDP-43-MBP-His_6_ with CK1δ + ATP (one out of two representative experiments is shown).

**Fig. S3.**
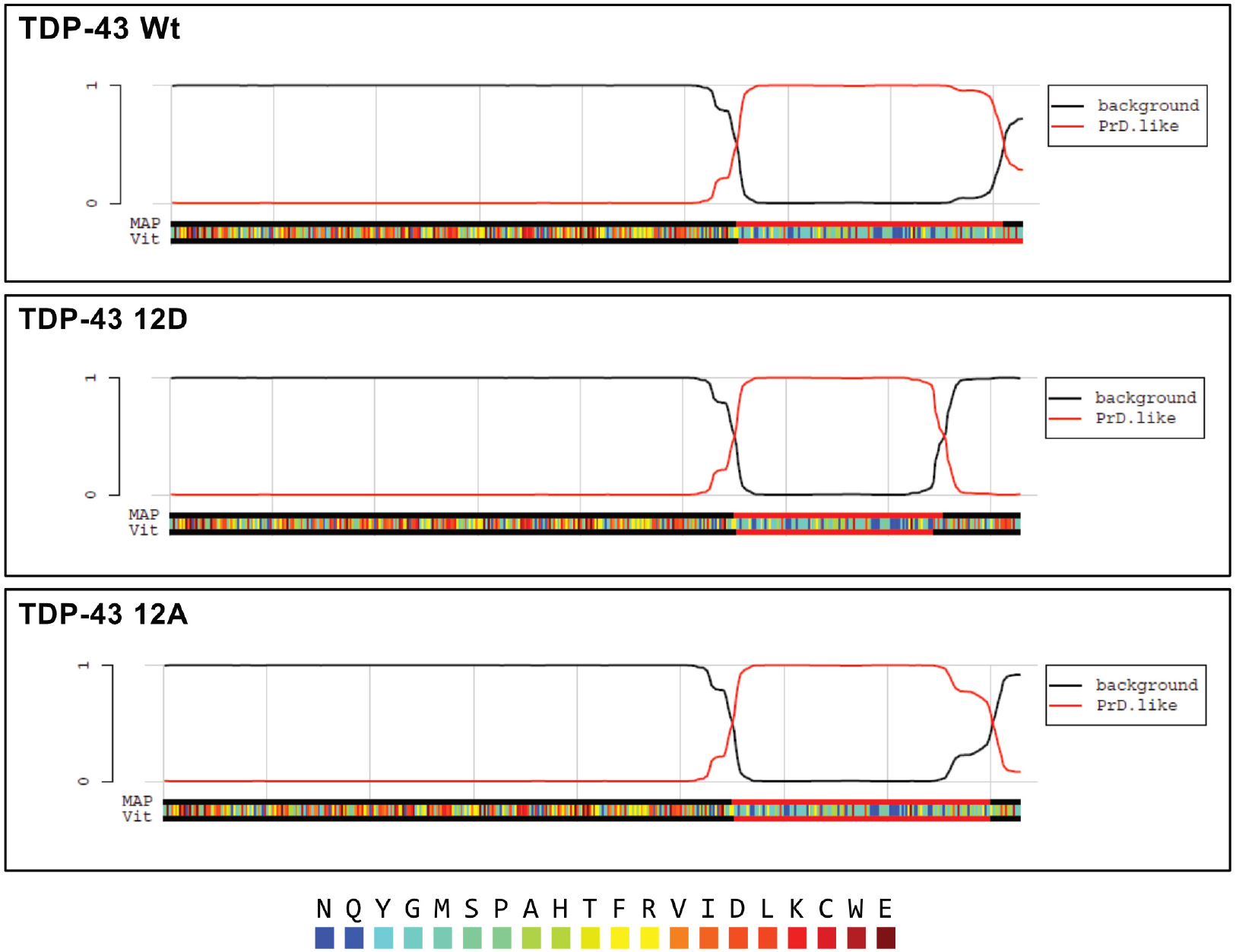
PLAAC predicts different prion sub-sequences on TDP-43 Wt vs. phosphomimetic TDP-43 12D. Visualization outputs from PLAAC, a web application that scans protein sequences for domains with **p**rion-**l**ike **a**mino **a**cid **c**omposition (Lancaster et al., 2014). Each box shows a detailed visualization of the TDP-43 protein variants (TDP-43 Wt, 12D and 12A) and respective Hidden Markov Model (HMM) prion-prediction score, showing that the prion-like character of the TDP-43 LCD is reduced by the 12 phosphomimetic S-to-D mutations. On the bottom, each amino acid is color-coded by its enrichment log-likelihood ratio in PrLDs with HMM parse indicated by outer bars (black and red).

**Fig. S4.**
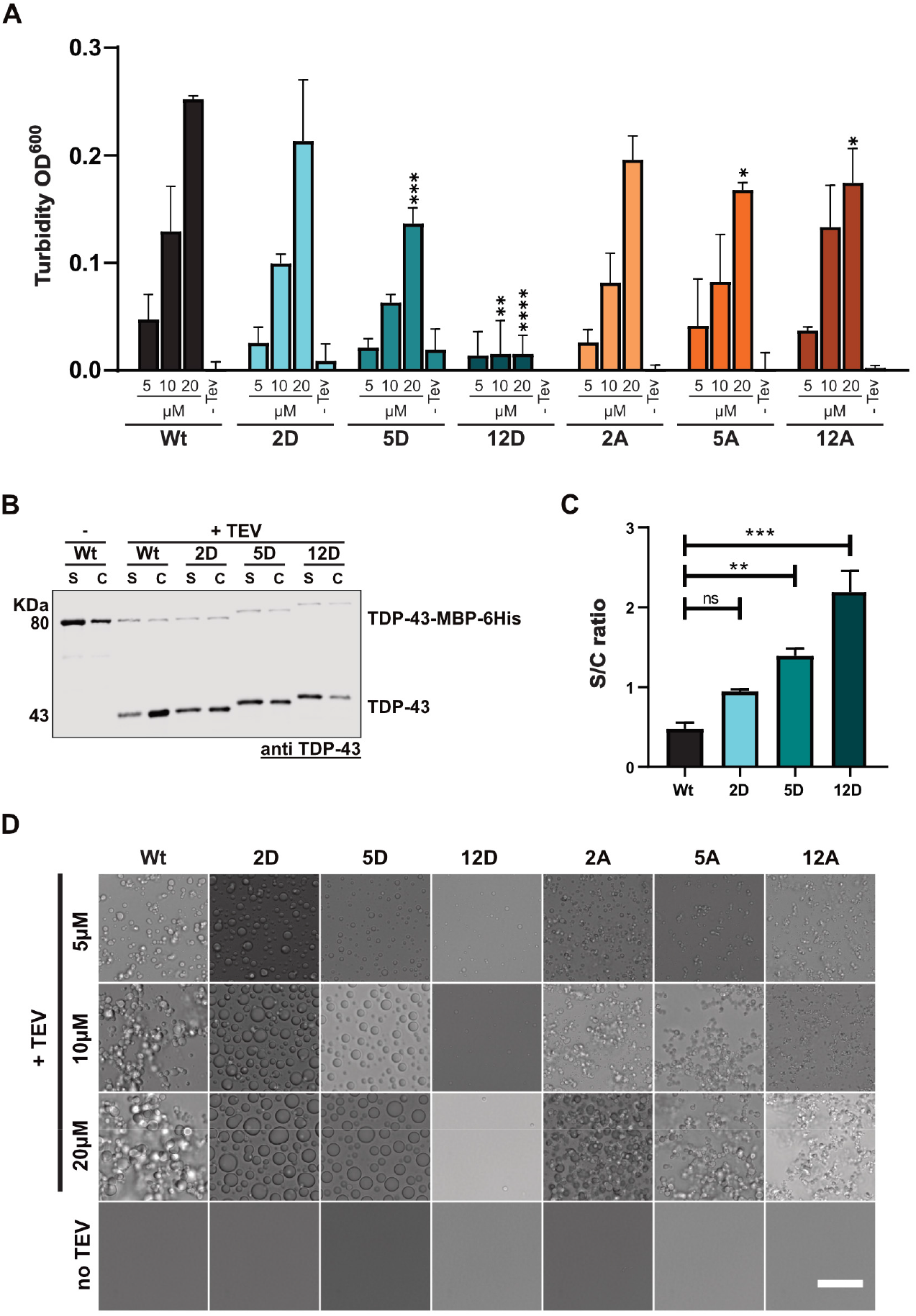
C-terminal phosphomimetic substitutions reduce TDP-43 condensation *in vitro*. A Turbidity measurements to quantify phase separation of different S-to-D and S-to-A mutants in comparison to TDP-43 Wt using phosphate buffer. Values represent means (n=3) ± SD. *p < 0.0332, **p < 0.0021, ***p < 0.0002 and ****p < 0.0001 by one-way ANOVA with Dunnett’s multiple comparison test to Wt. B Sedimentation assay to quantify condensation of S-to-D mutants in comparison to TDP-43 Wt (in Hepes buffer). TDP-43 was detected by TDP-43 Western blot (rabbit anti-TDP-43 N-term). C Quantification of band intensities corresponding to supernatant (S) and condensates (C) fractions is shown as means of S/C ratio (n=3) ± SEM. **p < 0.0021 and ***p < 0.0002 by one-way ANOVA with Dunnett’s multiple comparison test to Wt. D Representative bright field microscopic images of TDP-43 condensates in phosphate buffer (Bar, 25 μm).

**Fig. S5.**
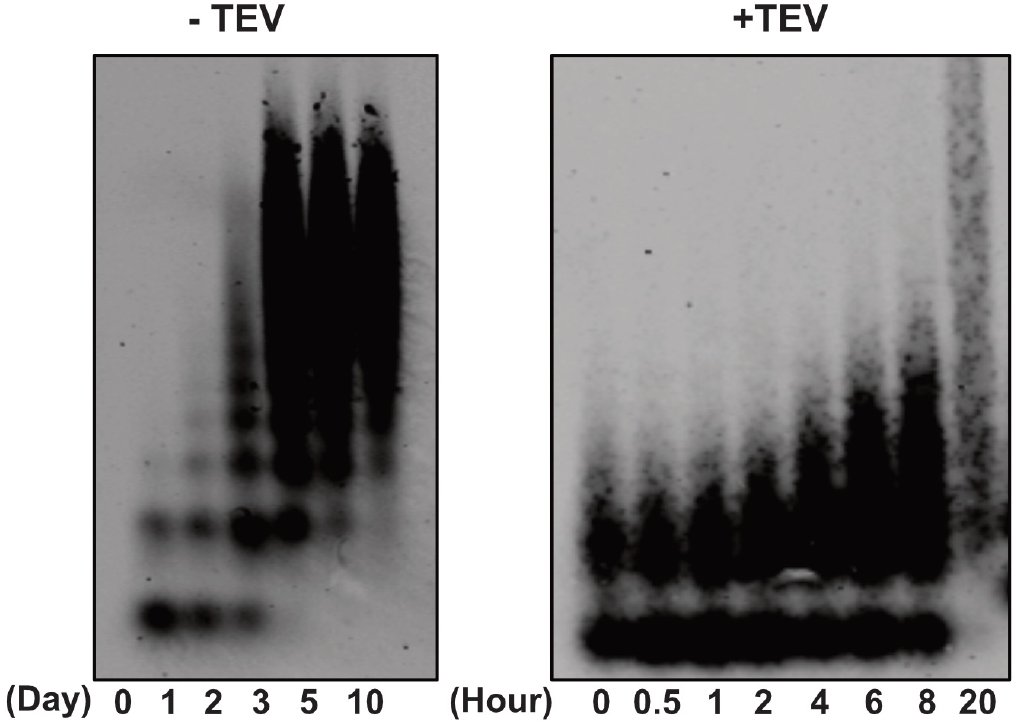
Comparison of SDD-AGE analysis of TDP-43-MBP-His_6_ with and without TEV protease-mediated cleavage. *In vitro* aggregation assay of TDP-43-MBP-His_6_ in the absence of TEV protease (-TEV) reveals the appearance of different SDS-resistant oligomers / high molecular weight species at late time points (at least 3 days of incubation). In comparison, when TDP-43-MBP-His_6_ is cleaved with TEV protease (+TEV), SDS-resistant oligomers / high molecular weight species already can be detected after several hours. Detection was performed after SDD-AGE analysis by anti-TDP-43 Western blot (rabbit anti-TDP-43 N-term).

**Fig. S6.**
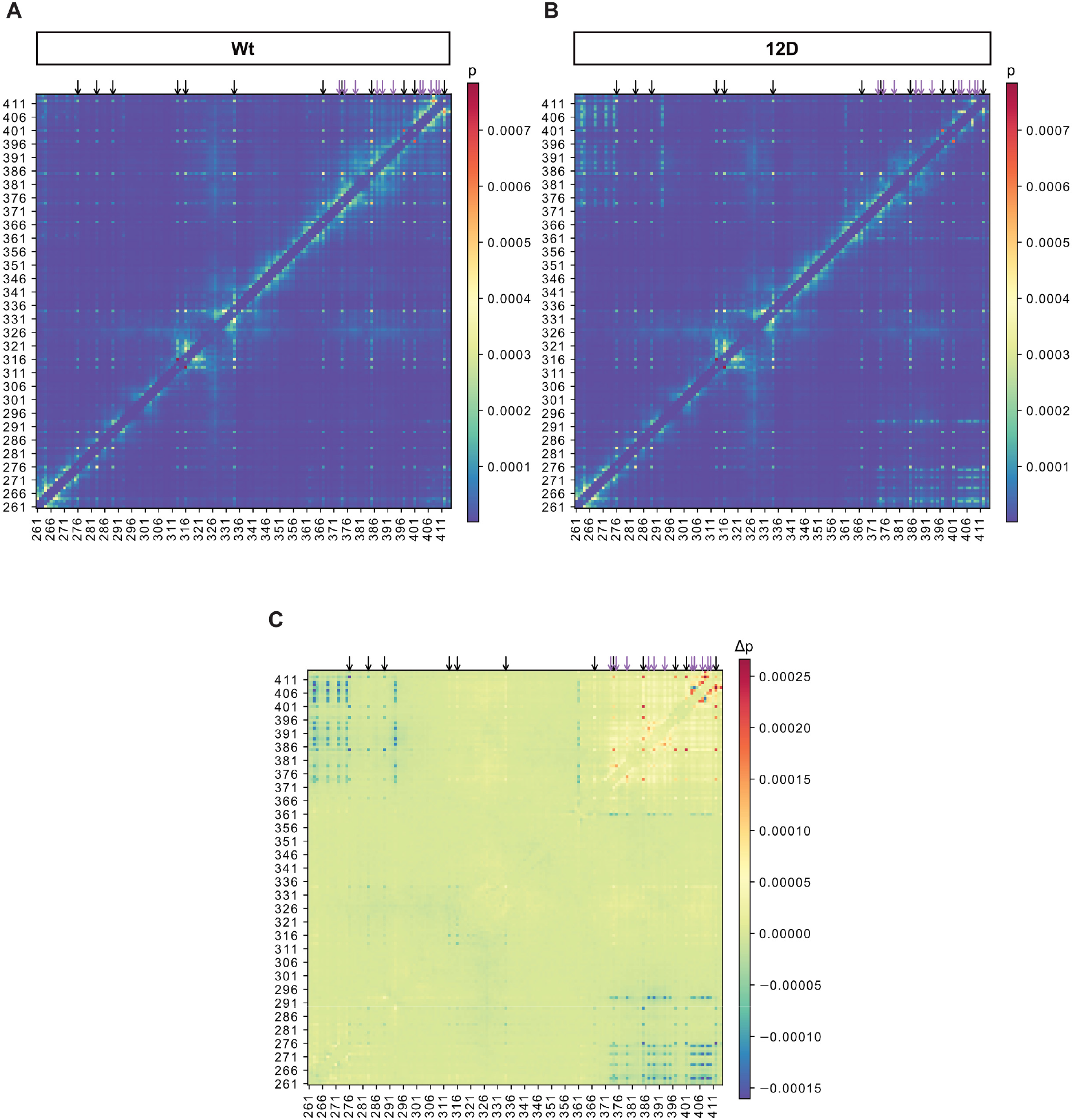
Analysis of contacts in biomolecular condensates formed by the TDP-43 LCD in coarse-grained simulations. A, B Contact maps for Wt (A) and 12D (B) TDP-43 LCD. Residue i and residue j are defined to be in contact if any of the coarse-grained beads are within 4.5 Å. The relative contact probability is calculated by averaging over all 118 protein chains and the last 5 μs of 20 μs simulations each. Intra-chain contacts with the two preceding and following residues are excluded from the analysis. Aromatic residues form prominent contacts and are highlighted by black arrows. E.g., looking at the column for F276 and following it upwards one can see that F276 interacts with F276 in other chains and irrespective of the chain, with F283, F289, F313, F316, W334, F367, Y347, W385, F401, and W412. The sites of the phosphomicking S-to-D mutations are highlighted by purple arrows. At these sites differences between Wt and 12D LCD can be seen, with Wt forming more contacts close in protein sequence and 12D instead interacting with R268, R272, R275, R293, and R361 further away in the sequence. C Differences in contact probability delta p_i,j = p_i,j(Wt) - p_i,j(12D) highlight that wild-type S residues, unlike phosphomicking D residues, favor interactions with residues close in sequence, while demonstrating that most contacts are not affected by the phosphomicking S-to-D mutations.

**Fig. S7.**
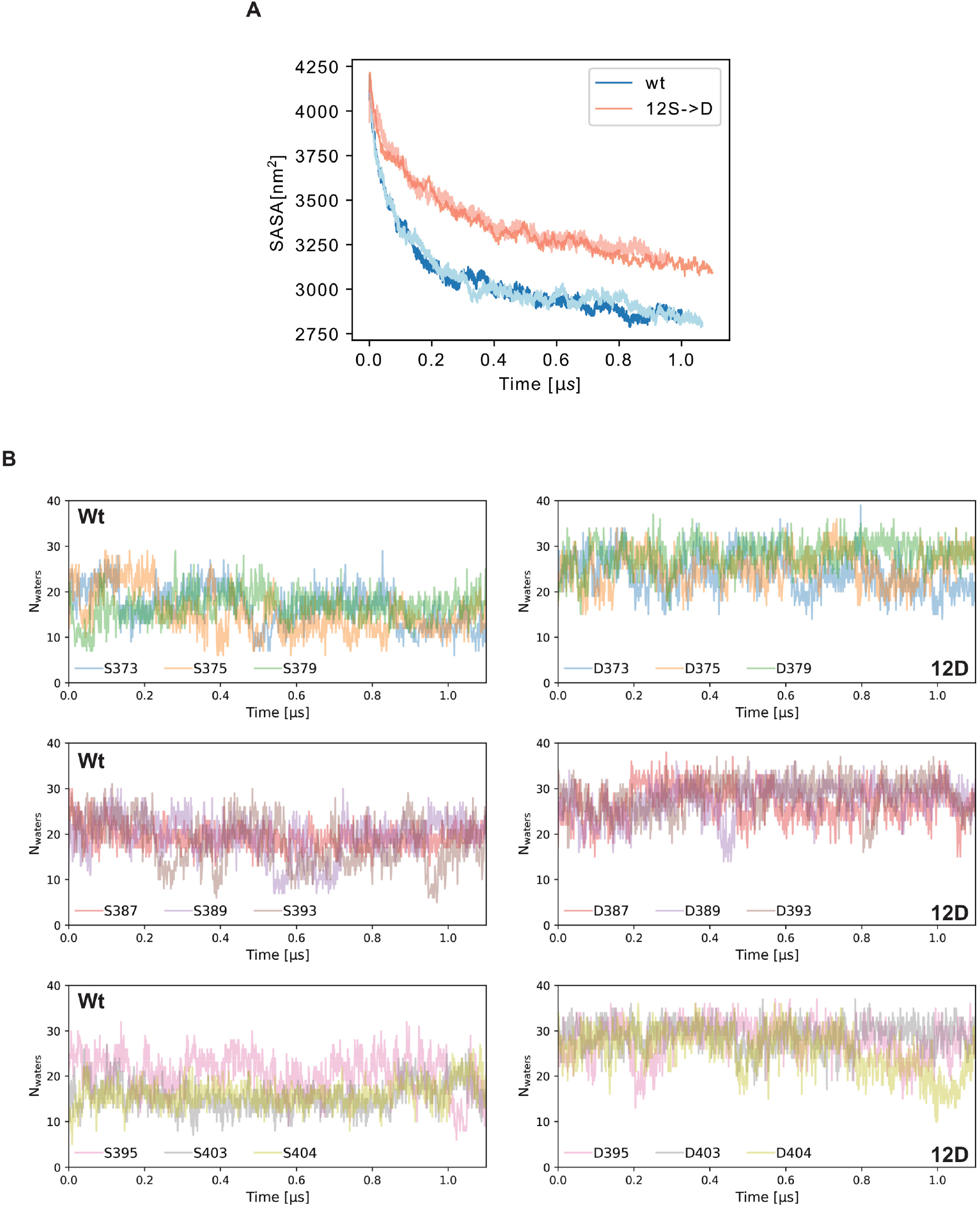
Differences in protein-protein and protein-water interactions in atomistic simulations of dense clusters of Wt and 12D TDP-43 LCDs. A Solvent-accessible surface area (SASA) of the LCDs of Wt (blue and light blue) and 12D mutant (red and salmon) in the simulations of their dense clusters as in biomolecular condensates. 12D mutant LCDs are more solvent-exposed than Wt LCDs, which is in line with the Wt LCDs forming stronger homotypic interactions. B Dynamics of sidechain-water interactions at the sites of phosphomimicking S->D substitutions in atomistic simulations of Wt and 12D LCD clusters. Numbers of waters bound to sites of phosphomimicking S-to-D mutations (within 5 Å of the sidechain) are tracked over time for Wt (left) and 12D LCDs (right). While interactions are dynamic, there is a consistent trend with the phosphomimicking Asp residues binding more water molecules than the Ser residues in Wt LCDs.

**Fig. S8.**
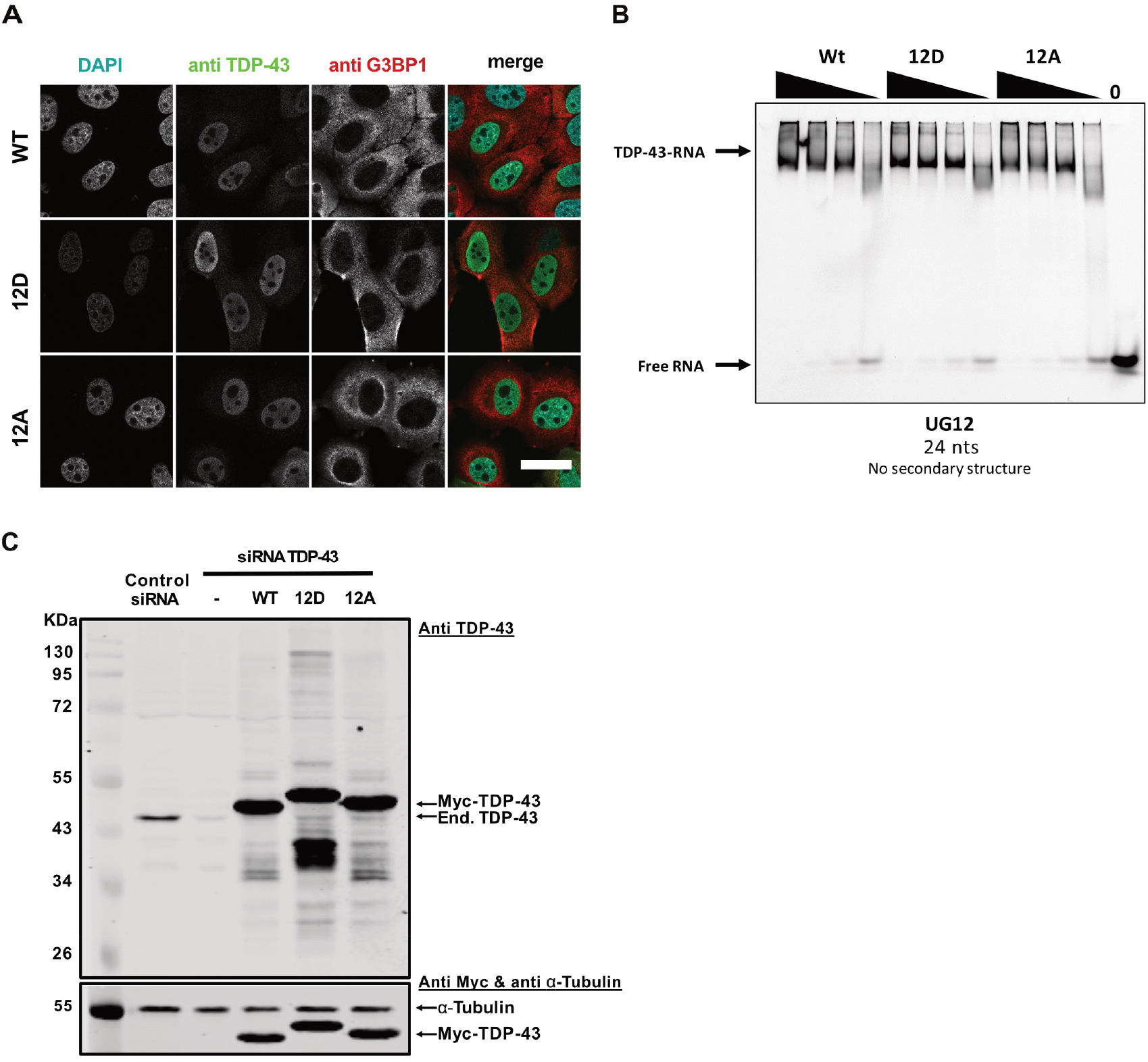
Phosphomimetic substitutions do not alter nuclear localization and UG-rich RNA binding of TDP-43. Control experiment for siRNA-mediated TDP-43 knockdown and reexpression of myc-tagged TDP-43 variants. A Immunostainings showing nuclear localization of TDP-43 Wt, 12D and 12A in HeLa cells. Endogenous TDP-43 expression was silenced by siRNAs, followed by transient transfection of the indicated siRNA-resistant myc-TDP-43 constructs. After 24 h, localization of TDP-43 Wt, 12D and 12A variants was visualized by TDP-43 immunostaining (mouse anti-TDP-43 antibody, Proteintech). G3BP1 (rabbit anti-G3BP1 antibody, Proteintech) and DAPI signal is shown to visualize the cytoplasm and nuclei, respectively. In the merge (right column), DAPI is show in turquoise, TDP-43 in green, and G3BP1 in magenta. Bar, 30 μm. B Electrophoretic mobility shift assay (EMSA) of TDP-43-MBP-His_6_ variants (Wt, 12D and 12A) in a complex with (UG)_12_ RNA. C SDS-PAGE followed by TDP-43 Western blot (upper blot) showing efficient siRNA-mediated knockdown of endogenous TDP-43 (running at ~43 kDa) in comparison to control siRNA and re-expression of myc-tagged TDP-43 Wt, 12D and 12A in Hela cells. Equal loading is demonstrated by α-Tubulin Western blot (bottom). TDP-43 was detected using rabbit anti-TDP-43 C-term antibody (Proteintech), α-Tubulin using mouse anti-alpha Tubulin antibody (Proteintech) and Myc-tag using mouse-anti Myc antibody (9E10, Helmholtz Center Munich).

**Fig. S9.**
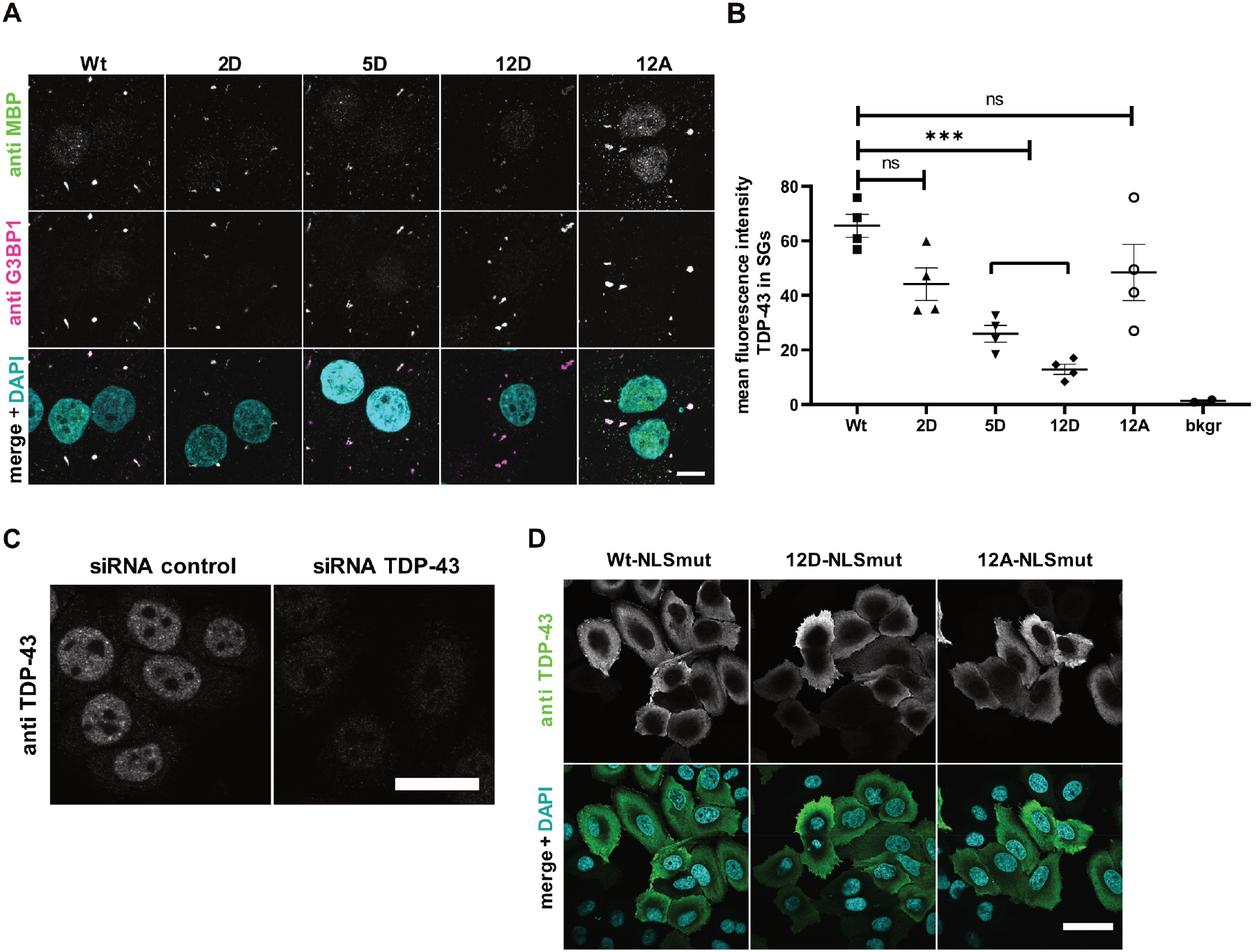
Phosphomimetic S-to-D substitutions reduce association of TDP-43 with stress granules in semi-permeabilized cells. Control experiments for siRNA-mediated knockdown and localization of TDP-43-NLS mutants. A Association of TDP-43 with stress granules (SGs) in semi-permeabilized HeLa cells is suppressed by phosphomimetic (2D, 5D and 12D) mutations in comparison to TDP-43 Wt and 12A. SGs and TDP-43-MBP-His_6_ were visualized by G3BP1 and MBP immunostaining, respectively. For clarity, signals were converted to grey values in the individual channels (upper two rows). In the merge (lower row), G3BP1 is shown in magenta, TDP-43-MBP-His_6_ in green, white pixels indicate colocalization. Nuclei were counterstained with DAPI (turquoise). Bar, 10 μm. B Quantification of the mean fluorescence intensity of TDP-43-MBP-His_6_ in SGs normalized to Wt for three independent replicates ± SEM, ***p < 0.0002 defined by 1-way ANOVA with Dunnett’s multiple comparison test (≥ 10 cells; ≥ 46SGs per condition). C Representative confocal images showing efficient siRNA-mediated knockdown of endogenous TDP-43 versus control siRNA transfection. Endogenous TDP-43 (grey) was detected using mouse anti-TDP-43 antibody (Proteintech). Bar, 25 μm. D Confocal images demonstrating equal cytoplasmic localization of TDP-43-NLS mutants (NLSmut Wt, 12D and 12A) 24 h after transfection and 48 h after endogenous TDP-43 silencing. Staining was carried out with mouse anti-TDP-43 antibody (Proteintech), signal shown in grey (upper row) or green (lower row); DAPI shown in turquoise. Bar, 40 μm.

**Movie 1.**

Fluorescently labelled TDP-43 Wt condensates imaged live by spinning disc confocal microcopy.

**Movie 2.**

Fluorescently labelled TDP-43 5D condensates imaged live by spinning disc confocal microcopy.

**Movie 3.**

Fluorescently labelled TDP-43 12D condensates imaged live by spinning disc confocal microcopy.

**Movie 4.**

Coarse-grained simulations of TDP-43 LCD Wt with explicit solvent.

**Movie 5.**

All-atom simulations of TDP-43 LCD 12D with explicit representation of proteins, ions, and water with atomic resolution.

